# A Gnotobiotic Rice Pathosystem Reveals How the Microbiome Suppresses Rice Blast Disease

**DOI:** 10.64898/2026.10.20.760547

**Authors:** Timothy R. Johnson, Lainey E. Kemmerer, Nalleli Payne, Matthew VanWeelden, Luiz F. Wurdig Roesch, Jessie Fernandez

## Abstract

Rice blast disease caused by the hemibiotrophic fungal pathogen *Magnaporthe oryzae* poses a major threat to global rice production, creating an urgent need for sustainable disease management strategies. Because plant-associated microbiomes can contribute to pathogen suppression and host defense, their role in rice blast resistance is of considerable interest. However, interactions among *M. oryzae*, rice, and the rice microbiome remain poorly understood. Using a gnotobiotic rice pathosystem, we investigated how microbiome colonization influences host responses to infection and how bacterial community structure changes following pathogen challenge. Microbiome colonization significantly reduced blast disease severity, while differential abundance analyses identified infection-associated shifts in bacterial community composition, including enrichment of *Brevibacillus*-associated taxa. Functional characterization of cultured endophytes revealed plant growth-promoting traits and antagonistic activity against *M. oryzae*. Two synthetic communities assembled from healthy- and infected-derived isolates significantly reduced disease severity, demonstrating that disease suppression can be recapitulated using simplified microbial communities. Interestingly, axenically grown plants exhibited elevated basal expression of multiple defense-associated genes but greater disease severity and reduced chitin-induced ROS production compared with holoxenic plants. In contrast, microbiome-colonized plants displayed enhanced ROS responsiveness and reduced disease, suggesting that the rice microbiome promotes a more effective and regulated immune state rather than simply increasing defense activation. Together, these findings advance our understanding of rice-microbiome-*M. oryzae* interactions and establish a tractable experimental platform for mechanistic studies of microbiome-mediated disease suppression.

**Highlights:**

- Gnotobiotic rice pathosystem enables controlled microbiome manipulation.
- Microbiome colonization significantly suppresses rice blast disease.
- Axenic plants show elevated defense-gene expression but greater disease.
- Rice microbiome colonization enhances chitin-induced ROS responses.
- Synthetic communities recapitulate microbiome-mediated disease suppression.

## 1. Introduction

Rice blast, caused by *Magnaporthe oryzae*, is one of the most economically important diseases of rice and remains difficult to control because of the pathogen’s highly effective infection strategy and adaptive capacity (Fernandez and Orth, 2018; Hu et al., 2022; Li et al., 2023, 2019). *M. oryzae* employs a hemibiotrophic lifestyle during infection, which begins with spore germination on the leaf surface and the formation of a specialized penetration structure known as an appressorium (Cruz-Mireles et al., 2021; Fernandez, 2023; Kalicharan and Fernandez, 2025). Once the leaf surface is penetrated by the appressorium, the fungus colonizes host tissue, initiating the biotrophic phase. Eventually, fungal colonization leads to host cell damage and the transition to the necrotrophic phase, accompanied by the development of characteristic blast lesions (Quime et al., 2025; Wilson and Talbot, 2009). Currently, disease management in the field is primarily achieved through fungicide application or the deployment of resistant cultivars; however, both strategies are costly, and pathogen populations can rapidly adapt to host resistance (Hu et al., 2022; Nalley et al., 2016). These limitations have increased interest in microbiome-based approaches that could complement existing methods of rice blast management. Although the molecular mechanisms underlying rice blast pathogenesis have been studied extensively, considerably less is known about how the rice-associated microbiome responds to infection or contributes to disease resistance.

Plant-associated microbial communities contribute to nutrient acquisition, development, stress tolerance, and protection against pathogens (de Andrade et al., 2023; Liu et al., 2021; Paasch and He, 2021). These functions may be supported by plant growth-promoting (PGP) bacteria through traits such as indole-3-acetic acid production (IAA), nitrogen fixation, phosphate solubilization, and direct antagonistic activity against pathogens. Disease suppression may result from direct microbial antagonism, competition for nutrients and colonization sites, production of antimicrobial metabolites, or modulation of host immune responses (Johnson et al., 2025; Kemmerer et al., 2026; Kour et al., 2024; Spence et al., 2014; Teixeira et al., 2021). Conversely, pathogen infection can alter the composition and functional potential of the host microbiome, potentially resulting in enrichment of microorganisms that either promote plant defense or contribute to disease progression (Seybold et al., 2020). Despite growing recognition of these interactions, it remains unclear how the rice microbiome responds to *M. oryzae* infection and whether infection-associated microbial taxa contribute functionally to rice blast resistance.

Plant immunity is shaped by pattern-triggered immunity (PTI), which is activated following recognition of conserved microbial or pathogen-associated molecular patterns and includes early responses such as the rapid production of reactive oxygen species (ROS) (Bigeard et al., 2015; Gao et al., 2021). Plant defense is also influenced by pathogen-induced systemic acquired resistance (SAR) and by beneficial microorganisms that can induce or prime systemic defense responses, commonly referred to as induced systemic resistance (ISR) (Gao et al., 2021; Hacquard et al., 2017). Although these mechanisms have been investigated extensively in dicot model systems, particularly *Arabidopsis*, important aspects of immune regulation differ in monocot crops such as rice (Balmer et al., 2013; Ke et al., 2019, 2014). These differences include the regulation and crosstalk of salicylic acid (SA), jasmonic acid (JA), and ethylene (ET) signaling pathways (Balmer et al., 2013; Vlot et al., 2021; Wan and Xin, 2022). Specifically, dicots exhibit minimal basal accumulation of SA but display rapid accumulation upon pathogen attack, which creates an antagonistic interaction with JA signaling (Wan and Xin, 2022). Conversely, monocots exhibit high basal SA concentrations, which are not significantly increased upon pathogen attack, and JA/SA crosstalk is not exclusively antagonistic, which is a prominent feature observed in rice immune response (Wan and Xin, 2022). Consequently, there remains a need for experimentally tractable systems that can resolve microbiome-host-pathogen interactions under controlled conditions.

Many studies examining the rice microbiome have relied on field-grown plants, where environmental heterogeneity can obscure mechanistic relationships between microbial community composition and disease outcomes (Dastogeer et al., 2022; Jobert et al., 2025; Tian et al., 2021). Gnotobiotic growth systems provide a complementary approach by enabling precise control of microbiome assembly while maintaining biologically relevant host-microbe interactions (Kremer et al., 2021; Vorholt et al., 2017). These systems have an advanced understanding of microbiome interactions in both dicot and monocot plants; however, controlled platforms integrating rice, *M. oryzae*, and the rice microbiome remain limited (Cheng et al., 2024; Kremer et al., 2021; Ma et al., 2022; Santos-Medellín et al., 2022).

Here, we developed a gnotobiotic rice pathosystem to investigate how the microbiome influences rice blast disease and how pathogen infection reshapes microbial community composition and function. Using this system, we compared immune responses in microbiome-colonized and axenic (microbe-free) plants, characterized infection-associated shifts in bacterial communities, evaluated cultured endophytes for PGP and antagonistic traits, and tested synthetic microbial communities for their ability to suppress disease. Together, our findings establish a controlled framework for dissecting rice-microbiome-pathogen interactions and show that microbiome colonization is associated with enhanced resistance to rice blast, altered host immune responses, and shifts in microbial community structure and functional potential.

## 2. Materials and methods

### 2.1. Strains and Media

The fungal strain *Magnaporthe oryzae* Guy11 was used for all experiments and maintained on complete medium (CM) agar (50 mL 20× nitrate salts, 1 mL trace elements, 10 g D-glucose, 2 g peptone, 1 g yeast extract, 1 g casamino acid, 1 mL vitamin solution, and distilled water to 1 L; pH 6.5; 15 g agar) at 24°C under a 12 h light/dark cycle. Endophytic bacteria were cultured on Reasoner’s 2A agar (R2A; Thermo Fisher Scientific, Waltham, MA, USA), tryptic soy agar (TSA; Thermo Fisher Scientific, Waltham, MA, USA), King’s B agar (KB; Millipore Sigma, Burlington, MA, USA), nutrient agar (NA; 5 g peptone, 3 g beef extract, 5 g NaCl, and distilled water to 1 L; pH 6.8; 15 g agar), or rice extract agar (RE). Rice extract agar contained 100 g four-week-old rice shoot tissue, 0.5 g yeast extract, and 1 g NaCl per liter. Rice tissue was boiled in 500 mL distilled water for 30 min, filtered, adjusted to 1 L with distilled water, and solidified with 15 g agar (pH 6.5). Endophytic bacterial isolates were grown in liquid culture at 28°C with shaking at 220 rpm. Long-term storage was achieved in 20% glycerol stocks maintained at −80°C.

### 2.2. Rice gnotobiotic growth conditions

Gnotobiotic growth chambers were generated using TP5000+TPD5000 Microbox containers (SacO₂, Deinze, Belgium). Calcined clay (AllSport Turface; SiteOne Landscape Supply, Gainesville, FL, USA) was used as a largely inert growth substrate to provide physical support while minimizing background microbial and nutritional inputs. Prior to assembly, the clay was thoroughly rinsed with DI water to remove fine particulates and autoclaved separately from the microboxes for 45 min. After cooling, sterile microboxes and clay were transferred to a biosafety cabinet for aseptic assembly. Each microbox received approximately 500 g of sterile clay and 200 mL of sterile 0.5× Murashige and Skoog (MS) liquid medium to support plant growth. The assembled gnotoboxes were subjected to a second 45 min autoclave cycle to ensure sterility prior to planting.

Dehusked rice seeds were surface sterilized (70% EtOH for 1 min, 2.5% bleach for 60 min, followed by five washes with sterile DI water) and aseptically sown into magenta boxes containing 0.5× MS medium solidified with Gelrite. Two days after sowing, seedlings were transferred to gnotoboxes. Five days after emergence, holoxenic (HO) plants received 2 mL of a soil inoculum prepared as a 50 g L⁻¹ soil slurry suspended in 0.5× MS medium. Soil was collected from rice-production fields in the Everglades Agricultural Area near Belle Glade, Florida (26.595394° N, −80.724098° W; 26.725180° N, −80.562292° W; 26.696683° N, −80.400598° W). Axenic (AX) plants received 2 mL sterile 0.5× MS medium. Gnotoboxes were maintained in a Percival growth chamber with a 12 h photoperiod, 80% relative humidity, and temperatures of 25°C during the day and 24°C at night.

### 2.3. Spray infection assay

Three days after microbiome inoculation, AX and HO plants were spray-inoculated with *M. oryzae* Guy11 conidia at 5 × 10⁴ spores mL⁻¹. Spores were harvested from 10-day-old CM cultures, filtered through Miracloth to remove mycelial fragments, suspended in 0.2% gelatin, and adjusted to the desired concentration using a hemocytometer.

Disease severity was assessed 5 days post-inoculation. For each treatment, one leaf from each of three plants per gnotobox was collected and imaged using an Epson Perfection V600 scanner. The experiment was repeated three times (*n* = 9 gnotoboxes per treatment; 27 leaves). Diseased leaf area was quantified in ImageJ as a percentage of total leaf area. Leaf measurements were averaged within each gnotobox, and the gnotobox was treated as the biological replicate.

### 2.4. DNA Extraction

Rice leaves were harvested 13 days post-transplantation (HO and AX) or 5 days post-infection (HOINF and AXINF). All leaves from each box were harvested (100-500 mg). Prior to DNA extraction, an enrichment procedure was used to reduce host DNA. Surface-sterilized leaves were cut into 5 mm segments and loaded into 2 mL tubes containing two 2 mm stainless-steel beads and 1.5 mL sterile 1× PBS. Samples were homogenized for 20 s at 4 m/s in a FastPrep-24 Tissue Homogenizer (MP Biomedicals, Solon, OH, USA). Homogenization conditions were selected to disrupt plant tissue while minimizing bacterial cell lysis. Samples were centrifuged at 800 × g for 5 min at 4°C, and the supernatant was transferred to a new tube. The supernatant was treated with DNase I (Qiagen RNase-Free DNase Set; Qiagen, Hilden, Germany) and incubated at 37°C for 30 min to deplete extracellular host DNA. EDTA was added to a final concentration of 10 mM to terminate the reaction. Intact bacterial cells were pelleted by centrifugation at 15,000 × g for 15 min at 4°C, washed with 1 mL cold PBS containing 10 mM EDTA, and subjected to DNA extraction using the Qiagen PowerLyzer PowerSoil DNA Extraction Kit according to the manufacturer’s instructions, beginning with resuspension in 750 µL PowerBead solution and 60 µL solution C1.

### 2.5. 16s rRNA amplicon sequencing and processing

DNA extracted from three independent experiments with three biological replicates per treatment (n = 9 per treatment) was used for targeted amplicon sequencing of the V3-V4 region of the bacterial 16S rRNA gene. DNA concentrations were quantified using the QuantiFluor dsDNA System and Quantus Fluorometer (Promega, Madison, WI, USA). Library preparation, quality control, and incorporation of peptide nucleic acid (PNA) blockers targeting the plant mitochondrial and chloroplast V3-V4 regions were performed prior to sequencing. Sequencing was conducted on an Illumina MiSeq platform (2 × 300 bp), and demultiplexed FASTQ files were returned. Sequencing adapters were removed using *Cutadapt* V5.2, and reads underwent quality-based filtering via the *DADA2* V1.30.0 pipeline (Callahan et al., 2016; Martin, 2011). Following denoising and chimera removal, the merged ASV table underwent taxonomic assignment. Taxonomy was assigned using the DADA2 implementation of the RDP naive Bayesian classifier with the SILVA v138.2 reference database (Quast et al., 2013). Reads assigned to mitochondria and chloroplasts were removed to generate a filtered ASV table.

### 2.6. Microbiome Data Analysis

The *Phyloseq* v1.46.0 package (McMurdie and Holmes, 2013) was used for microbiome analyses in R v4.3.2 (R Core Team, 2023). Sequencing yield and alpha diversity were initially compared between AX and HO samples using the filtered, unrarefied dataset. Samples with fewer than 1,000 non-organelle bacterial reads were excluded from HO versus HOINF community analyses, resulting in the removal of two HO and three HOINF samples. Alpha diversity was evaluated using observed ASVs and the Shannon diversity index. Because sequencing depth varied among samples, alpha-diversity analyses were performed on the rarefied datasets, with statistical comparisons conducted using linear models that included experimental batch as a covariate. Beta diversity was calculated from the filtered, unrarefied ASV table using Bray-Curtis and Jaccard dissimilarities. Differences in community structure were assessed by PERMANOVA, including batch as a covariate, and homogeneity of dispersion was evaluated prior to analysis (Oksanen et al., 2025). Ellipses were drawn to group sample types within a 95% confidence interval. PERMANOVA was performed using the adonis2 function in the *vegan* package, with the model distance ∼ Batch + Treatment, 9999 permutations, and marginal tests of terms using by = "margin". Homogeneity of variance was tested via the betadisper function from the *vegan* package.

Differential abundance analyses were performed using DESeq2 v1.42.1 (Love et al., 2014). Low-prevalence ASVs (<10 reads in fewer than two samples) were removed prior to analysis. ASV abundance was modeled as a function of batch and treatment using the Wald test with parametric dispersion estimation. Differentially abundant ASVs were identified using Benjamini-Hochberg adjusted *P*-values < 0.05. Heatmaps were generated from DESeq2-normalized counts summarized by treatment and transformed as log10(normalized counts + 1). All plots were made using the ggplot2 v3.5.2 package (Wickham, 2016).

### 2.7. Transcriptional Analysis of Rice Immune State via qPCR

Rice leaves were collected from plants at the same sampling time points used for microbiome analyses. Tissue was immediately frozen in liquid nitrogen and stored at −80°C until processing. Total RNA was extracted using the RNeasy Plant Mini Kit (Qiagen, Hilden, Germany) according to the manufacturer’s instructions and treated with DNase I to remove genomic DNA. RNA quality and concentration were assessed by spectrophotometry and agarose gel electrophoresis. First-strand cDNA was synthesized from 1 μg total RNA using a reverse-transcription kit with oligo-dT primers (QuantaBio, Beverly, MA, USA).

Quantitative PCR (qPCR) was performed using a SYBR Green-based detection system on a Bio-Rad real-time PCR instrument (Bio-Rad Laboratories, Hercules, CA, USA). Reactions were performed in technical quadruplicate using gene-specific primers (Table A-1). Target-gene expression was normalized to the rice housekeeping gene *GAPDH*, and relative expression was calculated using the ΔΔCT method. Expression values were reported relative to uninfected greenhouse-grown plants, which served as a nominal microbiome control. Three biological replicates were analyzed per treatment. Data visualization and statistical analyses were performed using GraphPad Prism v10.6.

### 2.8. ROS Burst Assay

ROS burst responses were measured in HO and AX plants as previously described with some minor modifications (Wang et al., 2022). Briefly, three plants per gnotobox were harvested, and their sheath tissue was cut into ∼3 mm segments with a sterile razor blade 1 day prior to the assay. Segments were placed in a 96-well plate containing 200 µL sterile ddH₂O and incubated overnight to attenuate wound responses. Tissue from each gnotobox was distributed across a single row, with two sheath segments per well.

ROS production was measured using a luminol-based assay. Solution A contained 0.6 mM luminol, 0.15 mM *p*-coumaric acid, and 8 µg mL⁻¹ horseradish peroxidase (HRP) in 50 mM Tris-HCl (pH 8.5), whereas Solution B contained 20 µM chitin CO6 oligomer (Elicityl, Crolles, France) in 50 mM Tris-HCl (pH 8.5). Water was replaced with 100 µL Solution A, and four baseline readings were recorded before addition of 100 µL Solution B to initiate elicitation. Luminescence was measured for 40 min at 56 s intervals with a 300 ms integration time. Traces were smoothed using a 3-point rolling average. Peak ROS production and total ROS production (area under the curve, AUC) were calculated from nine biological replicates per treatment.

### 2.9. Rice Leaf Endophyte Recovery

To recover and characterize rice leaf endophytes in the presence or absence of rice blast disease, leaves were harvested at the same sampling time points described above. Leaves were surface sterilized (10% EtOH, 1 min; 2.5% bleach, 5 min; five rinses with sterile water), and 80-100 mg of tissue was transferred to bead tubes containing 0.1 mm glass beads and two stainless steel beads. Samples were suspended in 1 mL 10 mM MgCl₂ and homogenized for 30 s at 4 m s⁻¹ using a FastPrep-24 homogenizer (MP Biomedicals, Solon, OH, USA). Homogenates were centrifuged at 1,000 × g for 1 min to remove large debris, and the resulting supernatant was serially diluted (10⁻¹-10⁻³) and plated on R2A, TSA, King’s B, nutrient agar, and rice extract agar. Representative colonies were purified and propagated in the corresponding liquid medium for long-term storage in 20% glycerol at −80°C. R2A medium was also used to validate microbiome colonization status in AX and HO plants.

### 2.10. MALDI-TOF Taxonomic Classification

To prepare isolates for MALDI-TOF analysis, isolates were first grown in liquid cultures in 0.5x TSB under the conditions outlined above. The cultures were grown overnight before being streaked onto 0.5x concentration TSA plates. The plates were incubated for 2-4 days at 28°C to acquire isolated colonies. Isolated colonies were picked and spotted onto the target slide. Sample preparation on the target slide was performed according to the manufacturer’s recommendations.

### 2.11. Plant Growth-Promoting Trait Assays

To evaluate plant growth-promoting traits, isolates were screened for indole-3-acetic acid (IAA) production, phosphate solubilization, and nitrogen-fixation activity. For all assays, bacterial cultures were prepared as described above and normalized to an OD₆₀₀ of 0.5 prior to analysis.

For IAA quantification, 3 mL cultures of 1/5× TSB supplemented with 0.5% (w/v) L-tryptophan were inoculated and incubated as described above. Following incubation, cultures were centrifuged at 10,000 rpm for 10 min, and 50 μL of supernatant was transferred to a 96-well plate. Salkowski reagent (50 mL 35% perchloric acid and 1 mL 0.5 M FeCl₃) was added (100 μL per well), and samples were incubated in the dark at room temperature for 30 min (Pramanik et al., 2023). Color development was visually assessed, with IAA-positive samples producing a pink coloration. Absorbance was measured at 536 nm using a microplate reader, and IAA concentrations were estimated using a standard curve.

Phosphate solubilization was assessed using Pikovskaya agar (PVK; 10 g D-glucose, 5 g Ca₃(PO₄)₂, 0.5 g (NH₄)₂SO₄, 0.2 g KCl, 0.1 g MgSO₄, trace amounts of MnSO₄ and FeSO₄, and 0.5 g yeast extract per liter) (Pramanik et al., 2023). A 5 μL aliquot of normalized culture was spotted onto PVK plates and incubated at 28°C for 5 days. Phosphate-solubilizing activity was quantified using the phosphate solubilization index (PSI):PSI = (halo diameter + colony diameter) / colony diameter Nitrogen-fixation activity was evaluated using nitrogen-free bromothymol blue (NFb) medium prepared with 5 g malic acid, 0.5 g K₂HPO₄, 0.2 g MgSO₄, 0.1 g NaCl, 0.02 g CaCl₂·2H₂O, 2 mL micronutrient solution (0.04 g CuSO₄·5H₂O, 0.12 g ZnSO₄·7H₂O, 1.40 g H₃BO₃, 1.0 g Na₂MoO₄·2H₂O, and 1.175 g MnSO₄·H₂O per liter), 2 mL bromothymol blue solution (5 g bromothymol blue in 0.2 M KOH), 4 mL Fe-EDTA solution (16.4 g Fe-EDTA per liter), 1 mL vitamin solution (10 mg biotin and 20 mg pyridoxine in 100 mL distilled water), and 4.5 g KOH per liter, with pH adjusted to 6.5 (Baldani et al., 2014; Cordova-Rodriguez et al., 2022). Normalized cultures were inoculated into NFb medium in a 96-well plate and incubated at 28°C for 24 h with shaking at 200 rpm. Absorbance was measured at 610 nm using a microplate reader. Color changes from green to blue were recorded as indicative of ammonium production, and fixed nitrogen concentrations were estimated using a standard curve (Cordova-Rodriguez et al., 2022).

### 2.12. *In vitro* Antagonism Assay

To assess antagonistic activity, overnight bacterial cultures were adjusted to an OD₆₀₀ of 0.5 with sterile water. A UV-sterilized 5 mm core borer, forceps, and rod were used to collect agar plugs from the actively growing edge of seven-day-old *M. oryzae* cultures. Fungal plugs were placed approximately in the center of a CM agar plate, ensuring mycelial contact with the media. On opposing sides of the fungal plug and equidistant from the center, two 5 µL droplets of normalized bacterial culture were spotted onto the plate. As a negative control, sterile water was applied in place of the bacterial inoculum. Plates were incubated at 25°C in the dark, and fungal growth was assessed at 5 days post-inoculation. Fungal colony diameters were measured and averaged for each replicate. The percentage of growth inhibition was calculated using the formula: Inhibition (%) = [(average control - average experimental)/average control] x 100. This applies to the diameter of the fungal colony.

### 2.13. Synthetic Community Formulation and Inoculation

To generate reproducible synthetic community inocula with defined concentrations of each isolate, OD600-to-CFU calibration curves were generated for each strain. Selected isolates were grown in 60 mL of 0.5× TSB, and 1 mL samples were collected every 2 h over an 8 h growth period. For each time point, OD600 was measured and serial dilutions (10^−3^ to 10^−6^) were plated in triplicate on 0.5× TSA. Following incubation at 28°C for 24 h, colony counts were recorded, log10-transformed, and plotted against the corresponding OD600 values to generate isolate-specific conversion equations relating OD600 to CFU mL^−1^ (CFU/mL = 10^(b^ ^+ m^ ^×^ ^OD^600^)^). These equations were used to estimate inoculum concentrations and determine the volume required to introduce approximately 200,000 cells of each isolate into each gnotobiotic system. Prior to inoculation, cells were pelleted and washed with 0.5× MS medium to remove residual growth medium. Synthetic community inocula were applied as a 2 mL suspension per gnotobox following the inoculation schedule described above.

### 2.14. Statistical Analysis

Statistical analyses were performed using GraphPad Prism 10.6 (GraphPad Software, Boston, MA, USA) and R. Disease severity was analyzed using one-way ANOVA followed by multiple-comparisons testing, with individual gnotobiotic systems treated as biological replicates. Comparisons of isolate functional traits (IAA production, nitrogen fixation, and phosphate-solubilization index [PSI]) between healthy- and infected-derived isolates were performed using Welch’s *t*-tests. Total ROS production was quantified as AUC, and peak ROS was defined as the maximum RLU observed during the assay. Differences in ROS responses were evaluated using two-tailed Welch’s *t*-tests. Microbiome analyses were conducted on filtered, unrarefied datasets after removing samples with fewer than 1,000 retained non-organelle bacterial reads. Retained read counts were analyzed using negative binomial generalized linear models (GLMs) with sequencing batch included as a covariate, whereas ASV richness was assessed using GLMs accounting for sequencing depth. Alpha-diversity metrics were analyzed using linear models that included treatment and sequencing batch as model terms. Differential abundance analyses were performed at the ASV level using DESeq2.

## 3. Results

### 3.1. Establishment and validation of a gnotobiotic rice growth system

To investigate microbiome-mediated effects on rice blast disease, we established a gnotobiotic rice growth system that enabled controlled manipulation of microbial colonization while maintaining normal plant development (Figure 1). Rice seedlings were inoculated with either a soil-derived microbial community to generate holoxenic (HO) plants or a sterile mock treatment to generate axenic (AX) plants.

**Figure 1.**
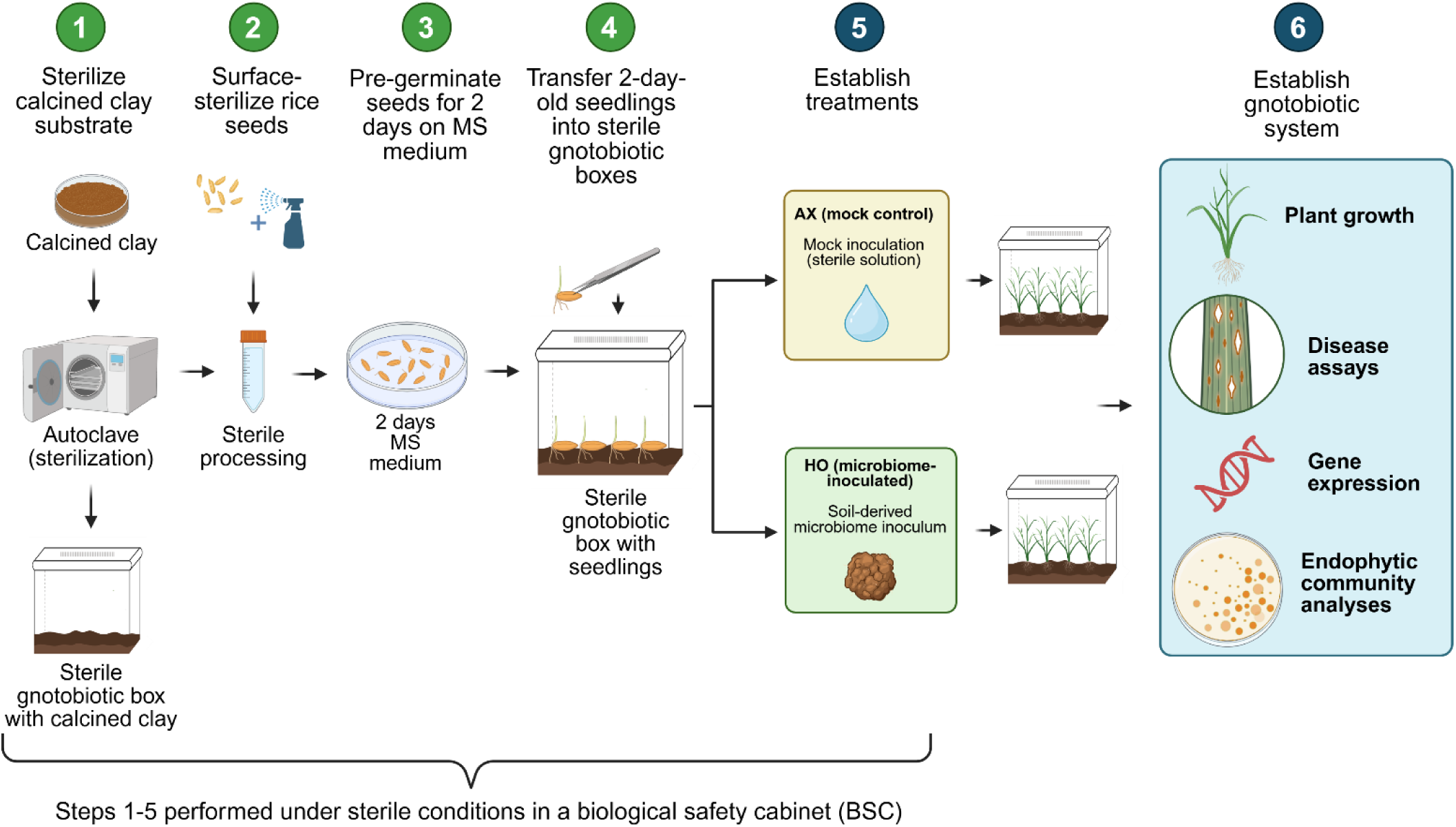
Diagram of the Gnotobiotic rice growth system. Illustration created in BioRender.com

To determine whether the system supported normal rice growth, plant development was compared between AX and HO conditions. No significant differences in plant growth were detected between treatments (Figures S1A-D). Plants grown under both conditions appeared healthy and exhibited comparable shoot and root development, with no obvious signs of nutrient deficiency or developmental abnormalities.

Microbial status was assessed using a culture-based recovery assay to confirm successful microbiome establishment in HO plants and effective microbial depletion in AX plants. Surface-sterilized leaf tissue from HO plants produced abundant microbial colonies when plated on R2A agar, whereas no detectable colonies were recovered from AX plants under the conditions tested (Figure S2E). These results were consistent across three independent experiments. Although the presence of low-abundance or unculturable microorganisms in AX plants cannot be entirely ruled out, the marked reduction in recoverable microorganisms indicates that the system maintained distinct microbial colonization states. Together, these results validate the gnotobiotic system as a platform for investigating microbiome-mediated effects on rice blast disease.

### 3.2. Microbiome colonization is associated with reduced rice blast severity

To assess the contribution of the rice microbiome to blast resistance, spray-infection assays using *M. oryzae* Guy11 were performed on HO and AX plants from two rice cultivars differing in blast susceptibility, the moderately resistant cultivar CO39 and the more susceptible cultivar YT16 (Figure 2 and Figure S2). For simplicity, infected HO and AX plants are hereafter referred to as HOINF and AXINF, respectively. Disease severity was quantified five days after inoculation as the percentage of diseased leaf area. Across both cultivars, AXINF plants consistently exhibited significantly greater disease severity than HOINF plants, suggesting that microbiome colonization contributes to blast resistance across distinct host genetic backgrounds. In CO39, HOINF plants exhibited an average diseased leaf area of 5.51%, whereas AXINF plants exhibited 28.15% diseased leaf area (*P* = 0.0043; Figure 2B). A similar pattern was observed in YT16, where HOINF plants displayed 3.6% diseased leaf area compared with 19.67% in AXINF plants (*P* = 0.0386; Figure S2B).

**Figure 2.**
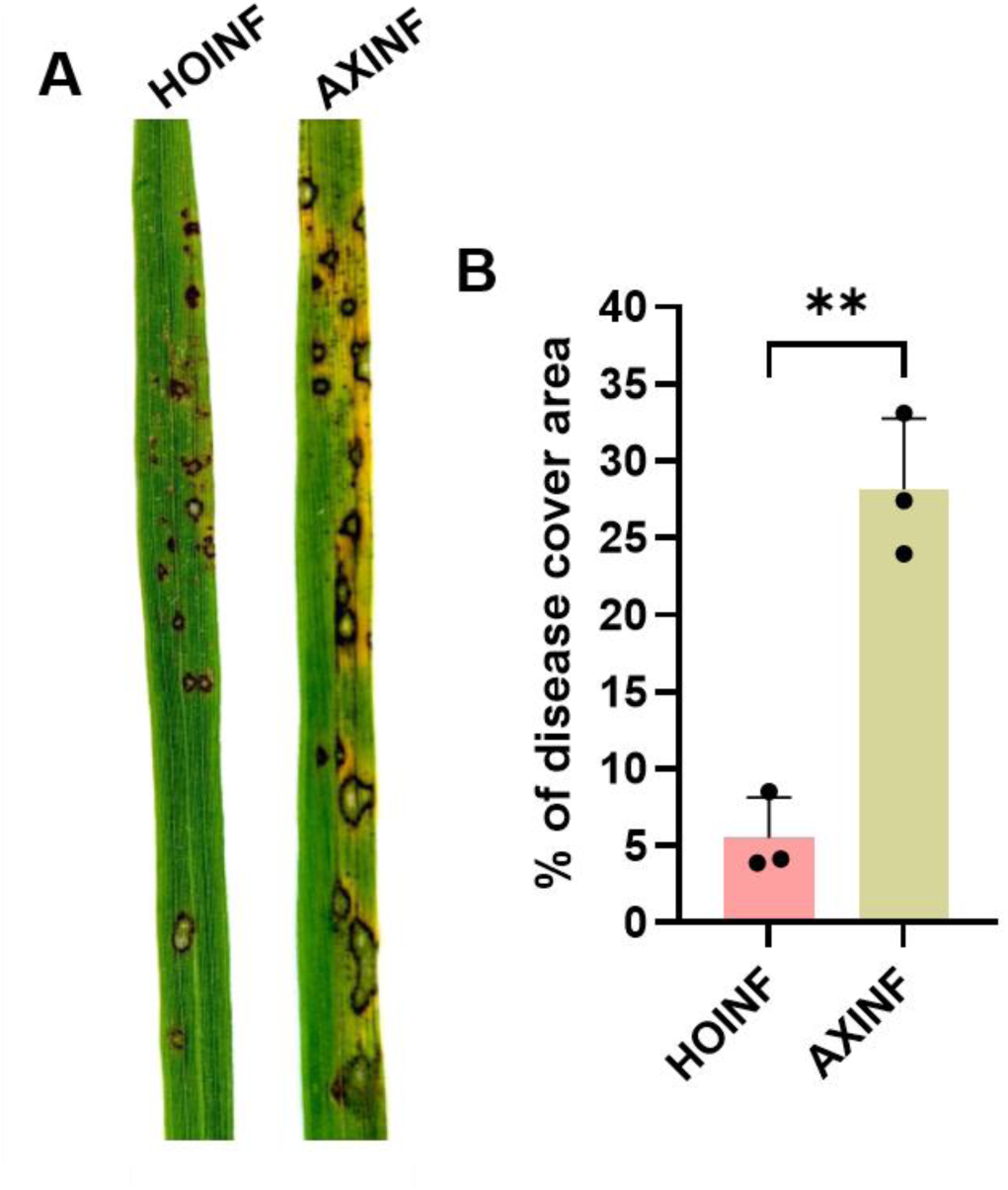
Holoxenic rice plants exhibit reduced blast disease severity compared with axenic plants. (A) Representative leaves of the rice cultivar CO39 showing blast symptoms following spray inoculation with *M. oryzae* (1 × 10⁴ spores mL⁻¹). (B) Disease severity is quantified as the percentage of disease-covered leaf area. Bars represent mean ± SD. Statistical significance was determined using Welch’s *t*-test. ** indicates *P* < 0.01. Holoxenic (HO); Axenic (AX)

In addition to quantitative differences in disease severity, clear differences in symptom development were observed between treatments. In both cultivars, AXINF plants exhibited more extensive yellowing surrounding lesions, indicative of increased disease-associated chlorosis (Figure 2A and Figure S2A). In contrast, chlorosis was reduced or absent in HOINF plants. Lesions on AXINF plants also appeared larger and more frequently coalesced than those observed on HOINF plants. Overall, these results demonstrate that microbiome colonization is associated with reduced rice blast severity and altered symptom development across multiple rice cultivars, supporting a role for the microbiome in suppressing disease progression during *M. oryzae* infection.

### 3.3. Axenic plants contain a reduced bacterial signal relative to holoxenic plants

To characterize the bacterial communities associated with gnotobiotically grown rice plants, 16S rRNA amplicon sequencing was performed on leaf endophytic samples. Raw and retained read counts for all treatments are summarized in Table S1. In addition to healthy and infected HO plants, AX plants were included to assess the level of bacterial signal recovered from culture-negative plants maintained under sterile growth conditions. AX plants yielded substantially fewer retained non-organelle bacterial reads than HO plants (Figure 3A; Table 1). Median retained read counts were 590 and 2,875 for AX and HO samples, respectively. Accordingly, AX samples fell below the 1,000-read threshold selected for downstream community analyses more frequently than HO samples, with 66.7% and 27.8% of samples failing to meet this threshold, respectively (Table 2). After accounting for sequencing batch effects, HO samples contained significantly more retained bacterial reads than AX samples, corresponding to an estimated 3.54-fold increase in bacterial signal (negative binomial GLM, Type II likelihood-ratio χ² = 4.45, *P* = 0.035; Figure 3A).

**Figure 3.**
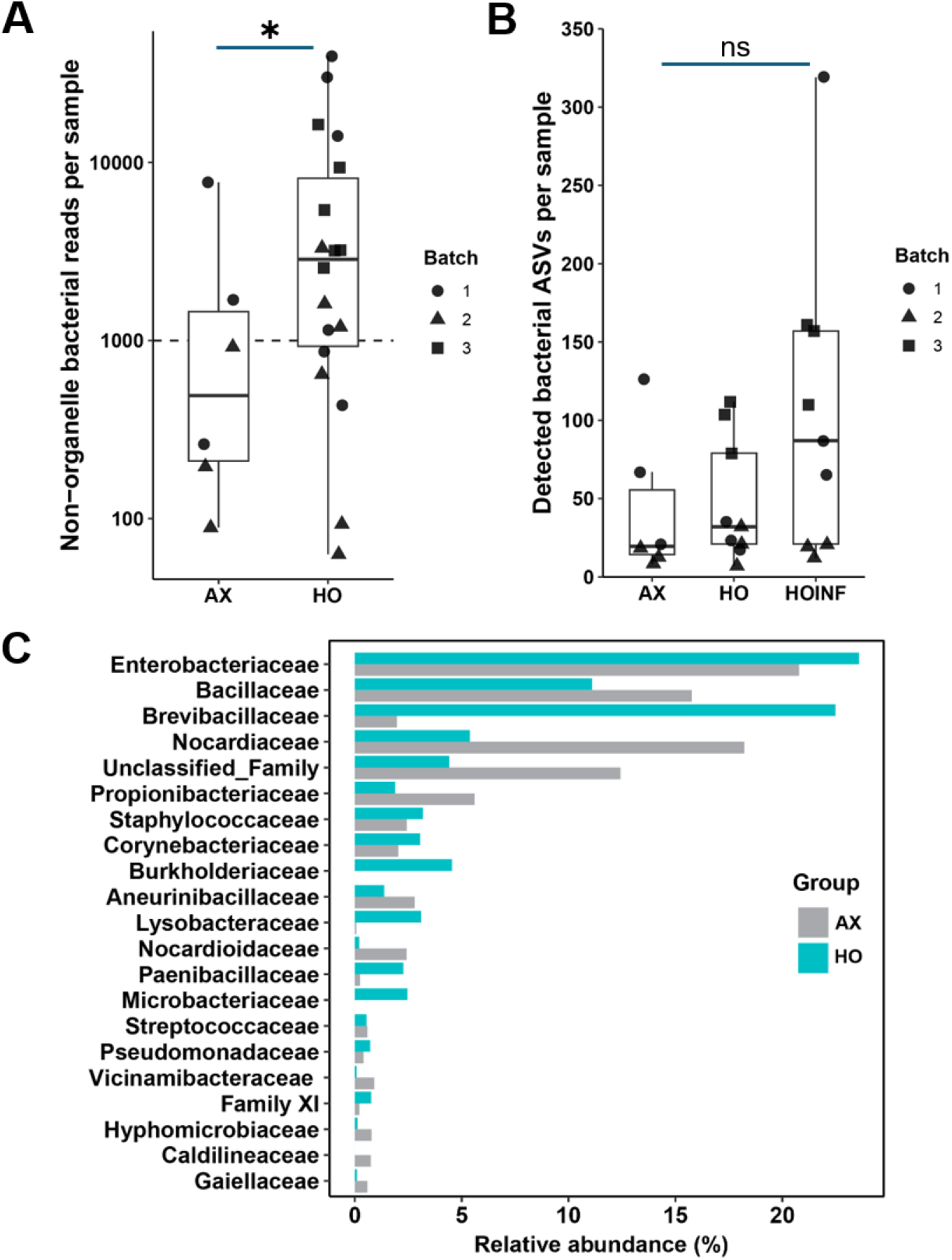
Microbiome composition and diversity of axenic and holoxenic rice plants. (A) Distribution of bacterial read counts assigned to AX and HO samples. The dashed line indicates the minimum sequencing depth threshold of 1,000 reads. Symbols denote independent experimental batches. (B) Distribution of detected amplicon sequence variants (ASVs) across AX, HO, and HOINF treatments. (C) Relative abundance of bacterial families detected in AX and HO/HOINF plants. Statistical significance in panel A was assessed using a negative binomial generalized linear model (GLM) with batch included as a covariate. Differences in panel B were evaluated using a generalized linear model with sequencing depth included as a covariate. * indicates P < 0.05; ns, not significant.

**Table 1.** Distribution of samples displaying less the 1000 retained reads per treatment type.

| Group | n | n Below 1000 Reads | Percent Below 1000 Reads |
| --- | --- | --- | --- |
| Axenic | 6 | 4 | 66.7% |
| Holoxenic | 18 | 5 | 27.8% |

**Table 2.** Summary of retained non-organelle bacterial read counts and detected amplicon sequence variants (ASVs) across treatment groups.

| Group | n | Median Reads | Mean Reads | Min Reads | Max Reads | Median Detected ASVs | Mean Detected ASVs |
| --- | --- | --- | --- | --- | --- | --- | --- |
| Axenic | 6 | 590 | 1815.67 | 89 | 7735 | 19.5 | 42.17 |
| Holoxenic | 18 | 2875 | 7395.06 | 63 | 39564 | 50 | 76.72 |

Consistent with the reduced bacterial signal, AX plants contained fewer detected amplicon sequence variants (ASVs) than HO plants, with median ASV counts of 19.5 and 50, respectively (Figure 3B; Table 1). However, after accounting for sequencing depth and batch effects, ASV richness did not differ significantly among treatments. Taxonomic profiles further highlighted differences between AX and HO plants (Figure 3C). Although bacterial reads were detected in AX samples, they were dominated by a few families, including Enterobacteriaceae, Nocardiaceae, Bacillaceae, and unclassified taxa. In contrast, HO plants exhibited greater representation of several plant-associated bacterial families, including Microbacteriaceae, Burkholderiaceae, Lysobacteriaceae, Paenibacillaceae, and Pseudomonadaceae (Figure 3C). These observations indicate that AX plants contained a substantially reduced bacterial signal relative to HO plants while remaining microbiologically distinct from the soil-derived communities detected in HO samples.

### 3.4. Rice blast infection alters endophytic bacterial community structure but not alpha diversity

Following quality filtering, HO and HOINF samples containing at least 1,000 retained non-organelle bacterial reads were included in downstream community analyses. HOINF samples exhibited modest increases in both observed ASV richness and Shannon diversity relative to HO samples; however, neither metric differed significantly between treatments after accounting for sequencing batch effects (Figure 4A). These results indicate that rice blast infection did not substantially alter within-sample bacterial diversity. Despite the limited changes in alpha diversity, differences in community composition were apparent at the family level (Figure 4B). HO communities were dominated by Enterobacteriaceae, with additional contributions from Brevibacillaceae and Bacillaceae. HOINF communities displayed a similar overall composition but were characterized by increased relative abundances of Brevibacillaceae and Bacillaceae and a corresponding reduction in Enterobacteriaceae.

**Figure 4.**
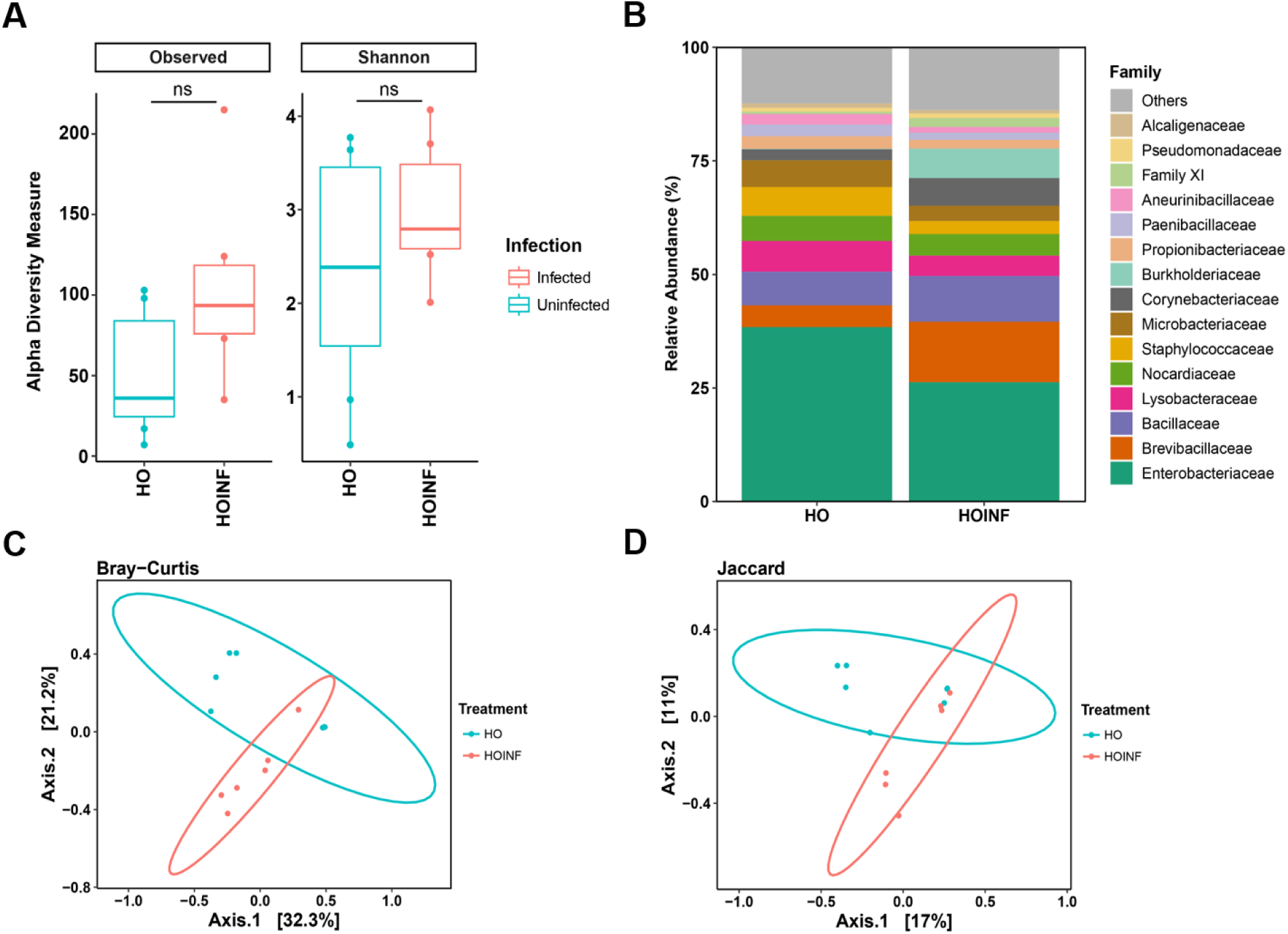
Bacterial community diversity, composition, and structure of HO and HOINF plants. (A) Alpha diversity metrics, including observed amplicon sequence variants (ASVs) and Shannon diversity index, for HO and HOINF samples. Points represent individual biological replicates (HO, *n* = 7; HOINF, *n* = 6), and horizontal bars indicate the mean ± SD. Differences in alpha diversity were evaluated using linear models with treatment and sequencing batch included as model terms; ns, not significant. (B) Relative abundance of the 15 most abundant bacterial families detected in HO and HOINF samples. Bars represent mean relative abundance across all biological replicates within each treatment. "Others" represents the combined abundance of all remaining low-abundance families. (C) Principal coordinate analysis (PCoA) based on Bray-Curtis dissimilarity showing abundance-weighted community structure. (D) PCoA based on Jaccard distance showing differences in community membership between HO and HOINF samples. Analyses were performed on the filtered, unrarefied dataset after removing samples with fewer than 1,000 non-organelle bacterial reads.

To determine whether infection altered overall community structure, Bray-Curtis and Jaccard distance metrics were analyzed using principal coordinate analysis (PCoA). Bray-Curtis ordination revealed partial separation between HO and HOINF communities (Figure 4C), and PERMANOVA identified a significant effect of infection on abundance-weighted community structure after accounting for sequencing batch effects (R² = 0.133, *P* = 0.015). In contrast, Jaccard-based analyses did not detect a significant effect of infection on community membership (Figure 4D; *P* = 0.188). These findings indicate that rice blast infection was associated primarily with shifts in the relative abundance of existing bacterial taxa rather than broad changes in community diversity or membership.

### 3.5. Rice blast infection is associated with enrichment of specific bacterial ASVs

Next, we sought to identify the bacterial taxa contributing to the infection-associated shifts in community structure observed in HOINF plants. Differential abundance analysis revealed several ASVs significantly associated with infection, with *Brevibacillus* (ASV25) showing the strongest enrichment in HOINF samples (Figure 5). Additional ASVs enriched in HOINF samples were assigned to the genera *Escherichia-Shigella*, *Corynebacterium*, *Rhodococcus*, and *Azospirillum*. In contrast, ASVs assigned to *Staphylococcus*, *Granulicatella*, *Rothia*, *Bradyrhizobium*, and *Lactobacillus* were more abundant in HO samples (Figure 5A).

**Figure 5.**
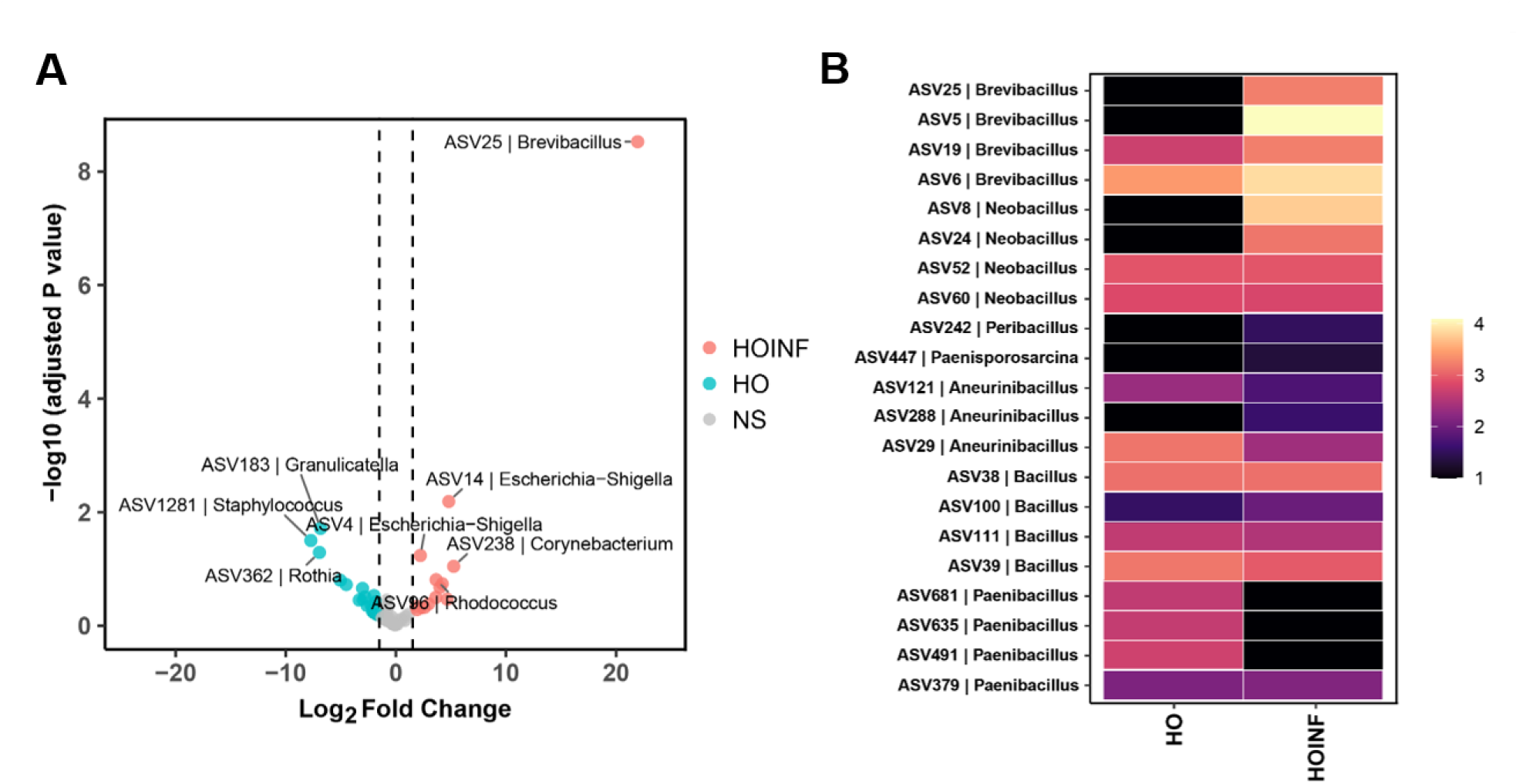
Differential abundance analysis of bacterial ASVs associated with HO and HOINF plants. (A) Volcano plot showing differential abundance of ASVs identified using DESeq2. Each point represents an individual ASV, and labeled points indicate genus-level taxonomic assignments when available. Log₂ fold change values were calculated relative to HO samples, such that positive values indicate enrichment in HOINF samples and negative values indicate enrichment in HO samples. (B) Heatmap of differentially abundant ASVs assigned to genera within the class *Bacilli*. Heatmaps depict DESeq2-normalized counts summarized by treatment and transformed as log10(normalized counts + 1). Differential abundance analysis was performed on the filtered, unrarefied dataset after removing samples with fewer than 1,000 non-organelle bacterial reads.

The enrichment of ASV25 was consistent with the increased relative abundance of Brevibacillaceae observed in HOINF communities (Figure 4B). However, infection-associated responses were not uniform within taxonomic groups. Notably, ASVs assigned to *Pseudomonas* exhibited both positive and negative log₂ fold-change values, suggesting contrasting responses among closely related taxa (Figure 5A). To further examine these patterns, normalized abundance heatmaps were generated for ASVs assigned to *Bacilli* and *Pseudomonas* (Figure 5B and Figure S3). Several *Brevibacillus* and *Neobacillus* ASVs displayed greater abundance in HOINF samples, whereas multiple *Paenibacillus*-associated ASVs were more abundant in HO samples (Figure 5B). Similarly, *Pseudomonas*-associated ASVs exhibited divergent responses to infection. ASV691, ASV1102, and particularly ASV159 were enriched in HOINF samples, whereas ASV605, ASV774, and ASV1050 displayed greater abundance in HO samples (Figure S3). These observations indicate that infection-associated shifts were driven by specific ASVs rather than uniform responses across broader taxonomic groups.

### 3.6. Microbiome-dependent differences in host immune signaling

To investigate how microbiome colonization influences host defense-associated gene expression during rice blast infection, we quantified the expression of genes associated with SA signaling, JA/ET signaling, phytoalexin biosynthesis, and pathogen-responsive defense pathways in HO, AX, HOINF, and AXINF plants (Figures 6 and 7). Although both YT16 and CO39 were initially evaluated, subsequent analyses focused on CO39 due to lower variability among biological replicates. Expression values are reported as fold change relative to age-matched greenhouse-grown plants.

**Figure 6.**
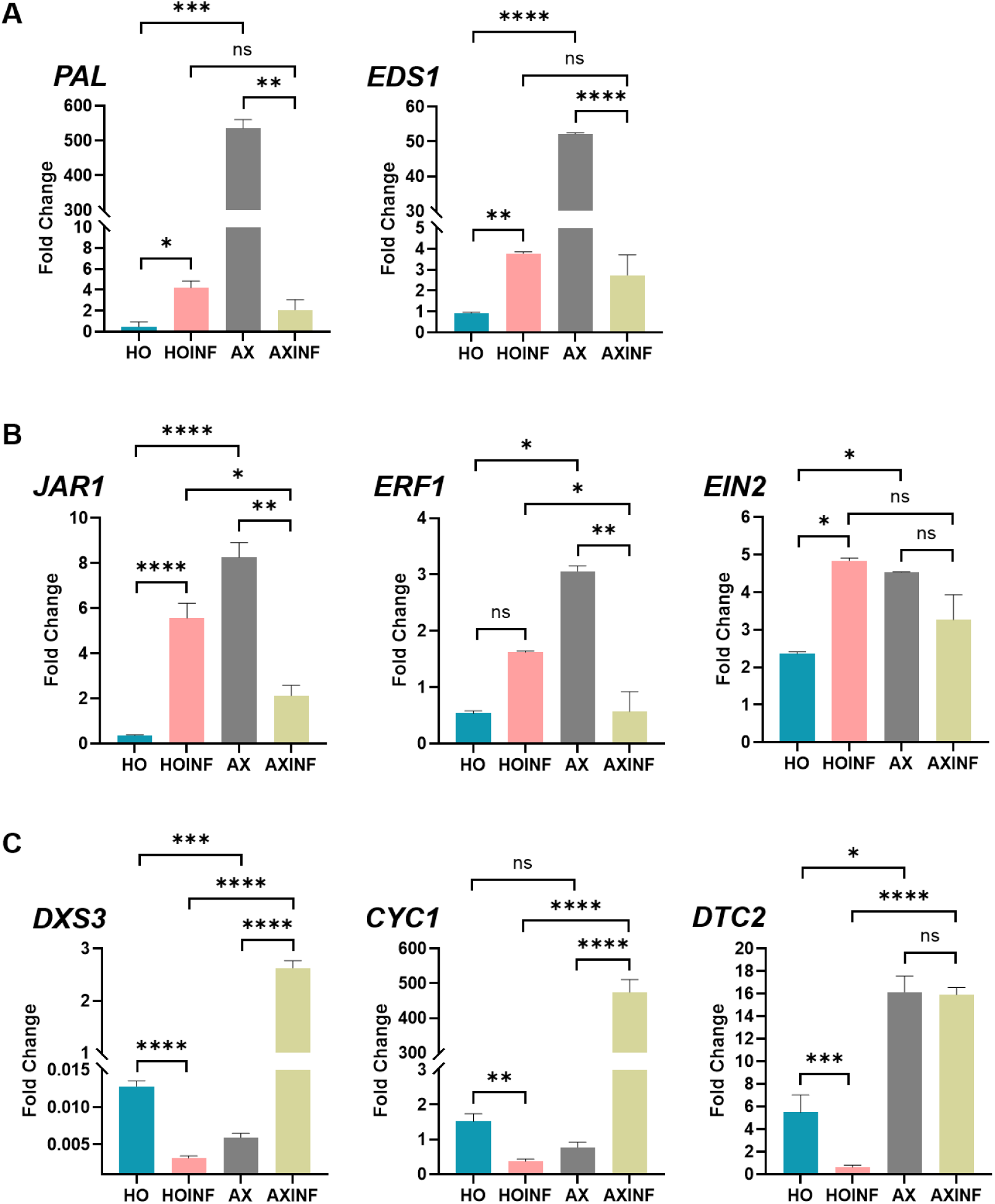
Effects of microbiome status on defense signaling and phytoalexin biosynthesis genes during rice blast infection. (A) Salicylic acid (SA)-responsive genes. (B) Jasmonic acid/ethylene (JA/ET)-responsive signaling genes. (C) Genes involved in the biosynthesis of the diterpenoid phytoalexin oryzalexin S. Gene expression is presented as fold change (2^−ΔΔCt^) relative to the calibrator control. Bars represent mean ± SD from three independent biological experiments. Statistical analyses were performed using ΔCt values and evaluated by two-way ANOVA followed by Šídák’s multiple-comparisons test. Asterisks indicate significant differences between treatments (*P* < 0.05, *P* < 0.01, *P* < 0.001, P < 0.0001); ns, not significant.

**Figure 7.**
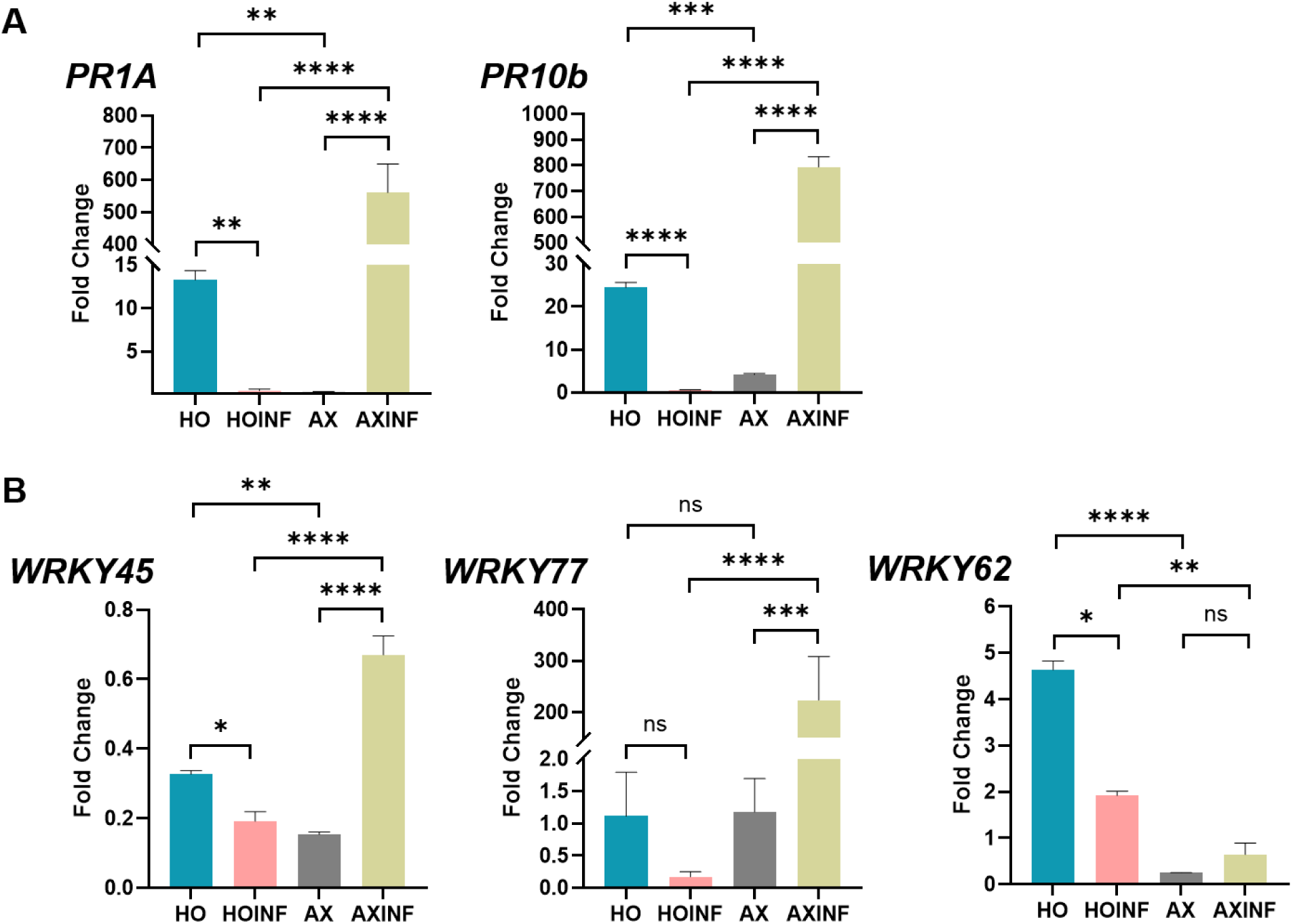
Effects of microbiome status on pathogen-responsive genes and WRKY transcription factors during rice blast infection. (A) Expression of pathogenesis-related (PR) genes. (B) Expression of WRKY transcription factors associated with plant immune responses. Gene expression is presented as fold change (2^−ΔΔCt^) relative to the calibrator control. Bars represent mean ± SD from three independent biological experiments. Statistical analyses were performed using ΔCt values and evaluated by two-way ANOVA followed by Šídák’s multiple-comparisons test. Asterisks indicate significant differences between treatments (*P* < 0.05, *P* < 0.01, *P* < 0.001, *P* < 0.0001); ns, not significant. HO, holoxenic; AX, axenic; HOINF, infected holoxenic; AXINF, infected axenic.

In the absence of infection, AX plants exhibited elevated expression of multiple defense-associated genes relative to HO plants. This response was observed across both SA- and JA/ET-associated pathways (Figure 6A, B). For example, *OsPAL* and *OsEDS1*, which are associated with SA biosynthesis and signaling, reached approximately 500-fold and 50-fold expression in AX plants, respectively, whereas expression remained near baseline levels in HO plants. Similarly, the JA/ET-associated genes *OsJAR1* and *OsEIN2* displayed increased expression in AX plants, with *OsEIN2* also exhibiting moderate upregulation in HO plants (Figure 6B). These results indicate that AX plants exhibited elevated basal expression of multiple defense-associated genes compared with microbiome-colonized plants.

Following infection, AXINF plants displayed a distinct transcriptional profile. Several genes that exhibited elevated expression in AX plants prior to infection, including *OsPAL*, *OsEDS1*, *OsJAR1*, and *OsEIN2*, were reduced following pathogen challenge (Figure 6A, B). In contrast, genes involved in diterpenoid phytoalexin biosynthesis were strongly induced in AXINF plants. *OsDXS3*, *OsCyc1*, and *OsDTC2*, which participate in the biosynthesis of the diterpenoid phytoalexin oryzalexin S (Valette et al., 2020a), exhibited substantial increases in expression in AXINF plants but showed little or no induction in HOINF plants (Figure 6C).

Marked differences were also observed among pathogen-responsive defense genes (Figure 7A). Expression of *OsPR1A* and *OsPR10B* was strongly induced in AXINF plants, reaching approximately 550-fold and 800-fold expression, respectively, whereas substantially lower expression levels were observed in the other treatments. Similarly, *OsWRKY77* reached approximately 150-fold expression in AXINF plants and displayed a transcriptional pattern closely resembling that of the PR genes (Figure 7B).

In contrast, WRKY transcription factors did not respond uniformly to infection (Figure 7B). *OsWRKY45*, a transcription factor commonly associated with SA signaling, remained below baseline expression across all treatments. *OsWRKY62* displayed a distinct pattern, exhibiting higher expression in HO and HOINF plants than in AX and AXINF plants. Overall, microbiome status was associated with distinct defense-associated transcriptional profiles. Uninfected AX plants exhibited elevated basal expression of several SA- and JA/ET-responsive genes, whereas infection of AX plants was associated with strong induction of PR genes and genes involved in oryzalexin S biosynthesis. These transcriptional responses were substantially reduced or absent in microbiome-colonized plants, indicating that microbiome status influences multiple defense-associated pathways during rice blast infection.

### 3.7. Microbiome colonization enhances chitin-induced ROS production

To determine whether microbiome colonization influenced pattern-triggered immune responsiveness, ROS production was measured following chitin elicitation. Leaf tissue collected from HO and AX plants was treated with chitin, and ROS production was monitored over time using a luminol-based assay. Chitin treatment induced a rapid ROS burst in both HO and AX plants; however, the magnitude of the response differed between treatments (Figure 8A). HO plants exhibited a significantly greater peak ROS response than AX plants (*P* = 0.016; Figure 8B). The ROS signal increased more rapidly in HO plants and remained elevated relative to AX plants throughout much of the assay period (Figure 8A). Consistent with this pattern, total ROS production, quantified as the area under the curve, was significantly greater in HO plants than in AX plants (*P* = 0.038; Figure 8C). These results indicate that microbiome colonization is associated with enhanced responsiveness to chitin elicitation. Despite elevated basal expression of several defense-associated genes in AX plants (Figures 6 and 7), HO plants exhibited a stronger chitin-induced ROS burst, indicating distinct immune-response profiles between microbiome-colonized and AX plants.

**Figure 8.**
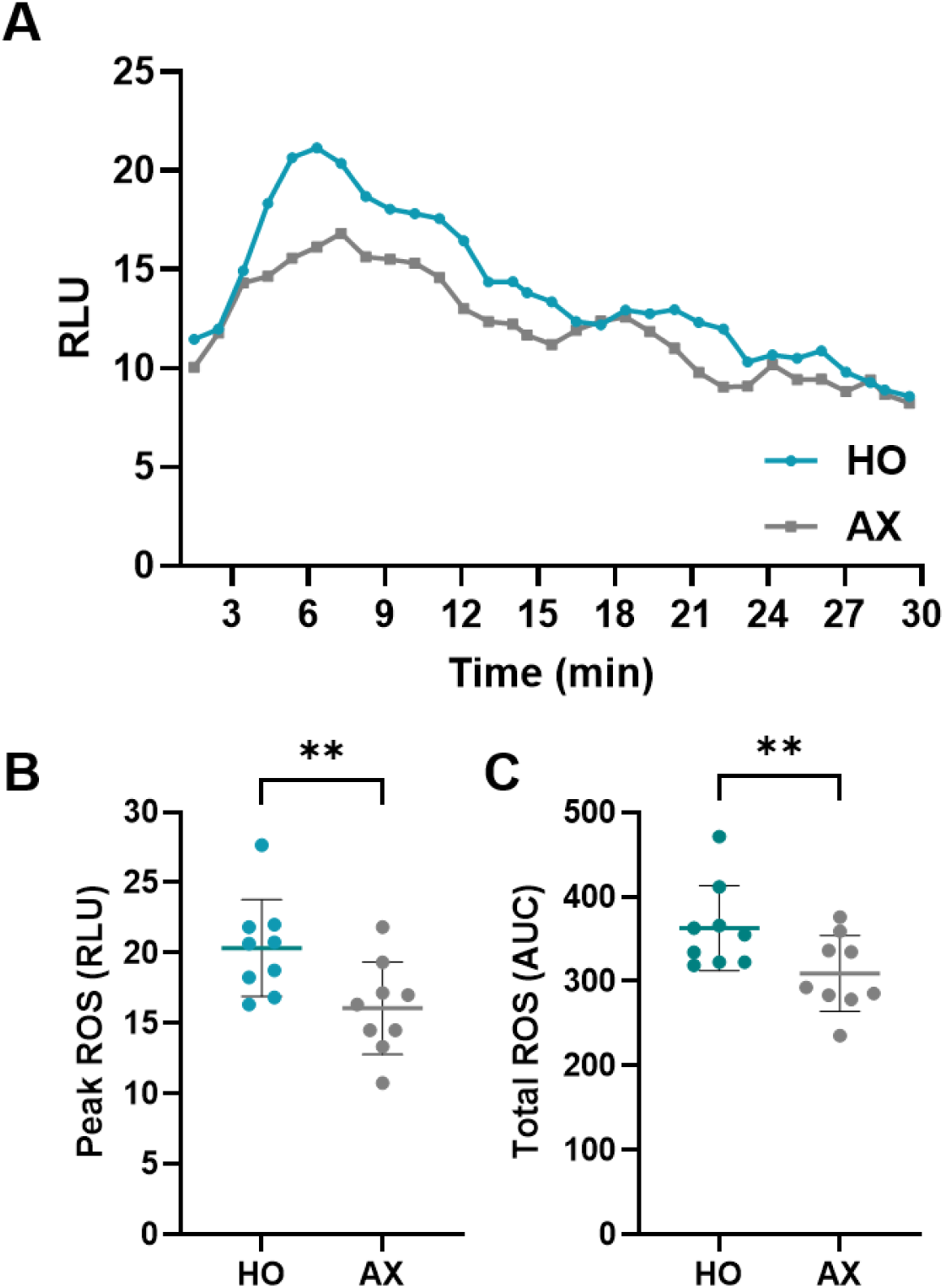
Microbiome status influences chitin-induced reactive oxygen species (ROS) production in rice. (A) ROS burst kinetics in HO and AX plants following treatment with 20 μM chitin oligomer over a 30-min period. Data represent mean relative luminescence units (RLU) from nine plants per treatment. (B) Peak ROS production is measured as the maximum RLU values. (C) Total ROS production was quantified by area under the curve (AUC) analysis. Points represent individual gnotoboxes, and horizontal bars indicate the mean. Data were obtained from three independent biological experiments comprising three gnotoboxes per treatment per experiment (*n* = 9 gnotoboxes per treatment). Statistical significance was determined using Welch’s *t*-test. ** indicates *P* < 0.01.

### 3.8. Functional characterization of microbiome isolates for SynCom development

In addition to culture-independent characterization by amplicon sequencing, culture-dependent analyses were performed to characterize cultivable members of the rice microbiome and support the construction of function-informed synthetic communities (SynComs). Following initial isolation and pre-screening, two sets of 10 isolates, representing healthy and infection-associated communities, were selected from a total of 47 recovered isolates for functional characterization and SynCom construction. Isolates were selected based on antagonistic activity, PGP traits, and MALDI-TOF-based taxonomic identification to retain a functionally diverse collection of strains.

Isolates recovered from HO plants represented a more taxonomically diverse group than those recovered from HOINF plants. Genera identified from HO plants included *Achromobacter*, *Cupriavidus*, *Pseudomonas*, and *Bacillus*. Although MALDI-TOF analysis produced species-level assignments that suggested diversity within these genera (e.g., *Achromobacter piechaudii* and *A. xylosoxidans*; *P. chlororaphis* and *P. stutzeri*), confidence scores did not meet the threshold required for reliable species-level identification (Table 3). Consequently, isolates are referred to at the genus level throughout this study. In contrast, isolates recovered from HOINF plants were less taxonomically diverse and consisted primarily of *Bacillus* and *Pseudomonas* representatives (Table 3).

**Table 3.** Endophytic bacterial isolates recovered from healthy and infected rice plants and their functional characteristics.

| ID | MALDI-TOF ID | % Inhibition | Phosphate |
| --- | --- | --- | --- |
| **Healthy** |  |  |  |
| RSC-H01* | *Pseudomonas usmongensis* | 15.1 ± 2.1 | N |
| RSC-H02 | *Cupriavidus necator* | 15.0 ± 3.1 | N |
| RSC-H03 | *Pseudomonas stutzeri* | 23.7 ± 9.8 | N |
| RSC-H04 | *Achromobacter xylosoxidans* | 18.4 ± 12.3 | N |
| RSC-H05 | *Achromobacter piechaudii* | 57.8 ± 26.4 | N |
| RSC-H06 | *Pseudomonas chlororaphis* | 59.0 ± 20.7 | Y |
| RSC-H07 | *Bacillus cereus* | 21.1 ± 6.8 | N |
| RSC-H08 | *Pseudomonas chlororaphis* | 39.7 ± 10.2 | N |
| RSC-H09 | *Pseudomonas stutzeri* | 25.6 ± 6.5 | N |
| RSC-H10 | *Pseudomonas chlororaphis* | 55.6 ± 16.8 | N |
| **Infected** |  |  |  |
| RSC-I01 | *Bacillus weihenstephanensis* | 52.7 ± 9.9 | N |
| RSC-I02 | *Pseudomonas usmongensis* | 5.3 ± 4.0 | Y |
| RSC-I03 | *Pseudomonas azotoformans* | 20.7 ± 4.8 | Y |
| RSC-I04 | *Pseudomonas azotoformans* | 21.2 ± 10.8 | Y |
| RSC-I05 | *Bacillus weihenstephanensis* | 55.7 ± 18.5 | N |
| RSC-I06 | *Pseudomonas usmongensis* | 71.3 ± 2.5 | Y |
| RSC-I07 | *Pseudomonas usmongensis* | 10.1 ± 3.4 | Y |
| RSC-I08 | *Pseudomonas usmongensis* | 11.1 ± 2.3 | Y |
| RSC-I09 | *Pseudomonas usmongensis* | 16.9 ± 7.5 | Y |
| RSC-I10 | *Bacillus weihenstephanensis* | 39.5 ± 11.2 | Y |

\*RSC-H, isolate recovered from a healthy plant; RSC-I, isolate recovered from an infected plant.

To compare PGP traits among isolate collections, we evaluated indole-3-acetic acid (IAA) production, nitrogen-fixation activity, phosphate solubilization, and antagonistic activity against *M. oryzae*. IAA production varied substantially among individual isolates but did not differ significantly between healthy- and infected-derived collections (Figure 9A, B; *P* = 0.739). Similarly, nitrogen-fixation activity exhibited considerable variation among isolates but did not differ significantly between groups (Figure 9C, D; *P* = 0.481).

**Figure 9.**
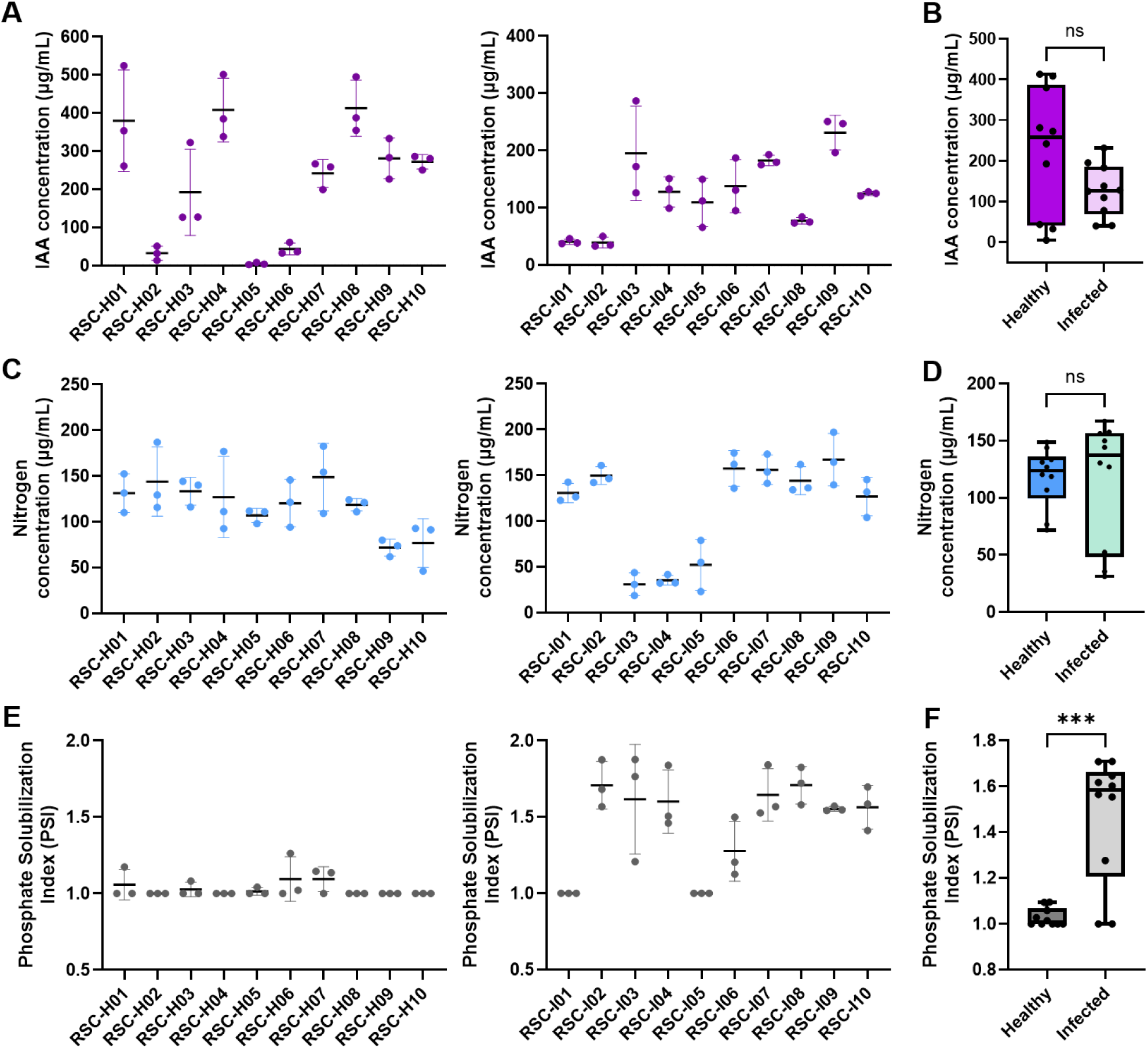
Antagonistic and plant growth-promoting traits of endophytic bacterial isolates recovered from healthy and infected plants. (A) Indole-3-acetic acid (IAA) production by individual isolates. (B) Comparison of mean IAA production between isolates recovered from healthy and infected plants. (C) Nitrogen fixation by individual isolates. (D) Comparison of mean nitrogen fixation between isolates recovered from healthy and infected plants. (E) Phosphate solubilization index (PSI) of individual isolates. PSI was calculated as the ratio of the total diameter of the colony and halo zone to colony diameter. (F) Comparison of PSI values between isolates recovered from healthy and infected plants. For panels A, B, D, and F, points represent measurements obtained from three independent experiments, and horizontal bars indicate mean ± SD. For panels C, E, and G, each point represents the mean value of an individual isolate calculated from three independent experiments. Statistical comparisons shown in panels C, E, and G were performed using Welch’s *t*-test. ***, *P* < 0.001; ns, not significant.

In contrast, phosphate-solubilization activity was enriched among infected-derived isolates (Figure 9E, F). PSI values were significantly greater in infected-derived isolates than in healthy-derived isolates (*P* < 0.001). Consistent with this pattern, eight of ten infected-derived isolates exhibited strong phosphate-solubilizing activity, whereas only one healthy-derived isolate consistently displayed this trait (Figure 9F; Table 3).

Antagonistic activity against *M. oryzae* was also observed among isolates from both collections (Table 3). Inhibition of fungal growth ranged from 5.3 to 71.3%. Several infection-derived isolates displayed strong antagonistic activity, including RSC-I01, RSC-I05, RSC-I06, and RSC-I10, all of which were assigned to either *Bacillus* or *Pseudomonas*. The strongest inhibition among infected-derived isolates was observed for RSC-I06 (*Pseudomonas* sp.), which inhibited fungal growth by 71.3 ± 2.5%. Among healthy-derived isolates, the strongest antagonistic activity was observed for RSC-H06 (*Pseudomonas* sp.) and RSC-H05 (*Achromobacter* sp.), which inhibited fungal growth by 59.0 ± 20.7% and 57.8 ± 26.4%, respectively. Collectively, these results indicate that antifungal activity was distributed across taxonomically diverse members of both isolate collections.

Overall, the selected isolates possessed diverse PGP and antifungal traits. While IAA production, nitrogen-fixation activity, and antagonistic activity were broadly distributed among isolates from both sources, phosphate-solubilizing activity was significantly enriched among isolates derived from infected plants. These functional characteristics provided the basis for subsequent SynCom construction and evaluation.

### 3.9. Reconstruction of protective communities using defined SynComs

To determine whether simplified microbial communities could reproduce the disease-suppressive effects observed in microbiome-colonized plants, isolates were grouped by source community and used to construct a Healthy SynCom (HSC) and an Infected SynCom (ISC) (Table 3). The two SynComs were designed to represent healthy- and infection-associated culturable communities identified through amplicon sequencing and culture-dependent analyses.

Representative disease symptoms are shown in Figure 10A. Plants inoculated with either SynCom developed fewer and smaller lesions than HOINF plants. Lesions in HSC- and ISC-treated plants were generally less extensive and exhibited reduced coalescence compared with those observed in HOINF plants. In addition, the chlorotic halos that were prominent in AXINF plants during the initial infection assays were largely absent in HOINF, HSC, and ISC plants (Figures 2A and 10A).

**Figure 10.**
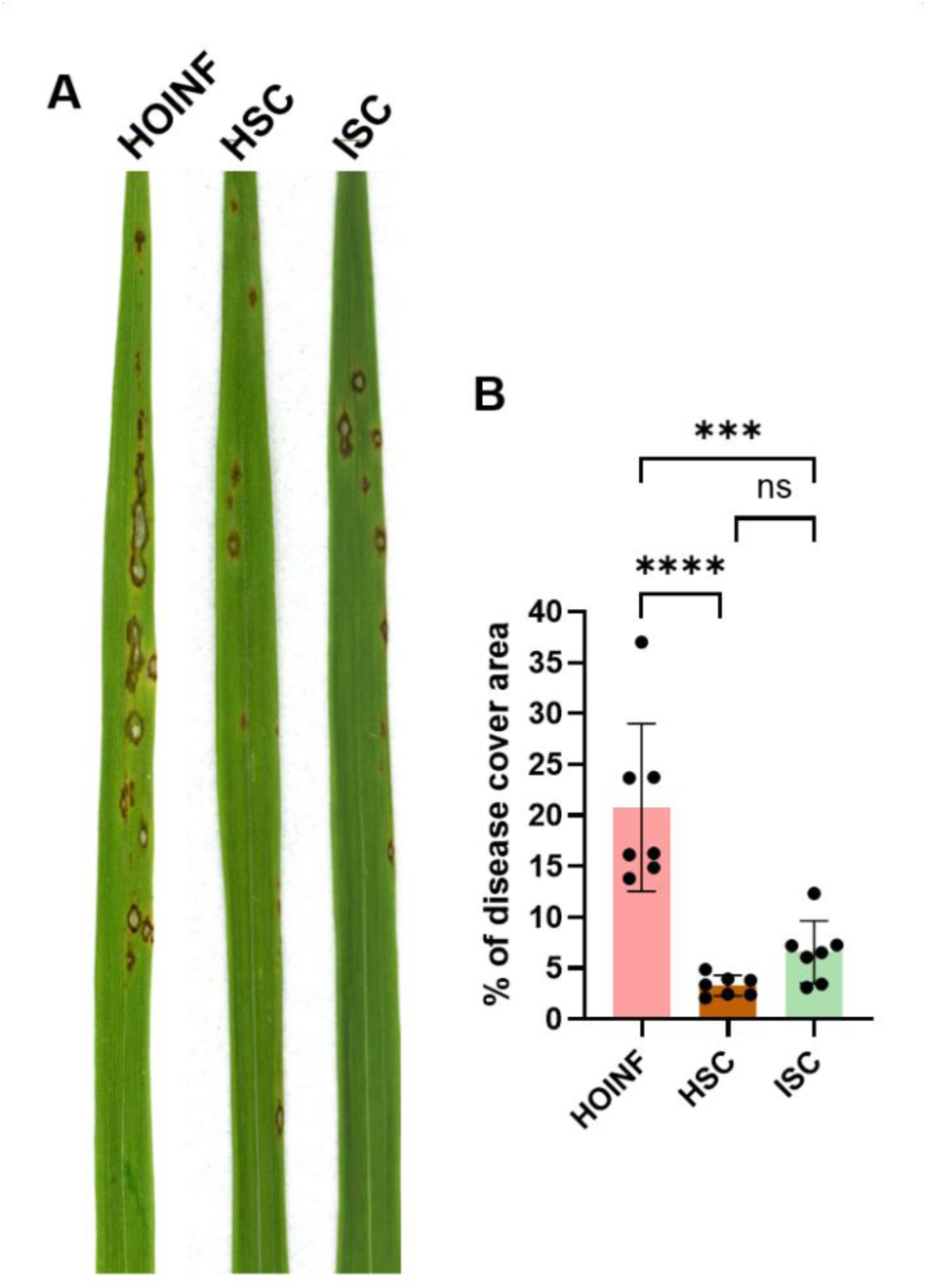
Effects of synthetic community (SynCom) inoculation on rice blast disease severity. (A) Representative leaves of HOINF, HSC, and ISC plants following inoculation with *M. oryzae*. (B) Disease severity is quantified as the percentage of disease-covered leaf area. Bars represent mean ± SD from three independent biological experiments. Statistical significance was determined using one-way ANOVA followed by Tukey’s multiple-comparisons testing. Asterisks indicate significant differences between treatments (***, *P* < 0.001; ****, *P* < 0.0001); ns, not significant. HOINF, infected holoxenic plants; HSC, healthy-derived synthetic community; ISC, infected-derived synthetic community.

**Figure 11.**
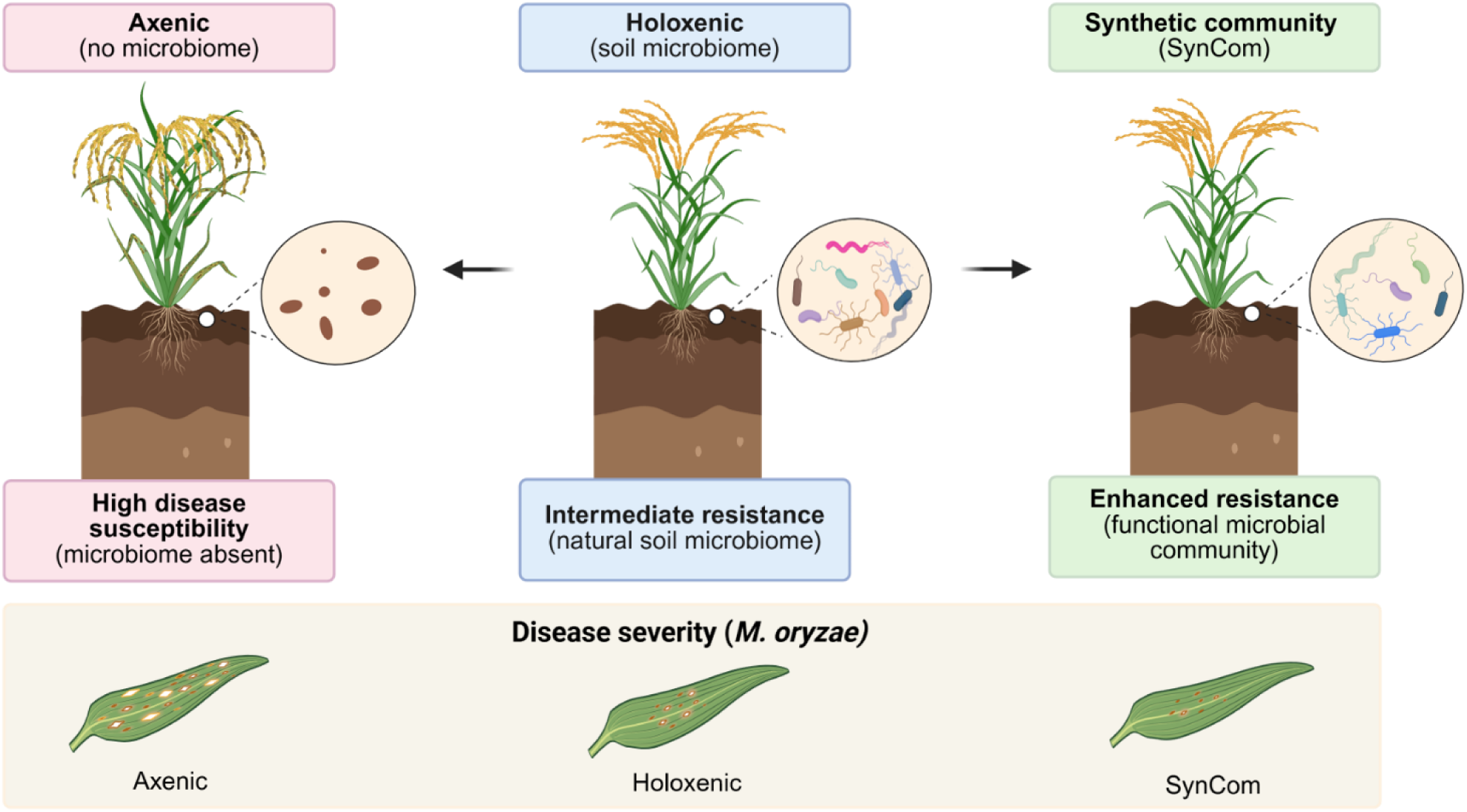
Microbiome colonization and synthetic communities suppress rice blast disease. Axenic rice plants lacking a microbiome exhibited the highest susceptibility to *M. oryzae*. Holoxenic plants colonized with a soil-derived microbiome displayed reduced disease severity, while synthetic communities (SynComs) reproduced much of the disease-suppressive effect observed in microbiome-colonized plants. Representative leaf phenotypes illustrate the relative disease severity observed in SynCom-treated, holoxenic, and axenic plants following *M. oryzae* infection.

These visual observations were supported by quantitative disease measurements (Figure 10B). Mean diseased leaf area was 20.91% in HOINF plants, compared with 3.97% and 6.07% in HSC- and ISC-treated plants, respectively. SynCom treatments significantly reduced disease severity relative to HOINF plants (both *P* < 0.0001). Although HSC-treated plants exhibited a lower mean diseased area than ISC-treated plants, the difference between the two SynCom treatments was not significant (*P* = 0.538). Despite the enrichment of phosphate-solubilizing isolates within the ISC (Figure 9F), SynCom origin did not significantly influence disease suppression. Both healthy- and infected-derived SynComs substantially reduced rice blast severity, indicating that simplified microbial communities are sufficient to reproduce much of the disease-suppressive effect observed in microbiome-colonized plants.

## 4. Discussion

Rice blast disease remains one of the most significant threats to rice production worldwide, creating a need for sustainable management strategies that reduce reliance on chemical inputs while maintaining crop productivity. Biological control and microbiome-based approaches represent promising alternatives, but the mechanisms through which the rice microbiome influences blast resistance remain poorly understood. Here, microbiome colonization reduced rice blast severity, altered host immune responses, and enabled the construction of simplified synthetic communities that also suppressed disease. Together, these findings demonstrate that the rice microbiome contributes to blast resistance and provide a controlled framework for investigating microbiome-mediated disease suppression.

Rice plants colonized with a complex soil-derived microbiome exhibited reduced lesion development compared with axenically grown plants. Axenic plants developed larger lesions with more extensive surrounding chlorosis, a phenotype consistent with a more susceptible interaction, whereas lesions on colonized plants remained comparatively restricted (Maina and Oerke, 2023). Microbiome-mediated resistance may arise through complementary mechanisms, including direct antagonism of the pathogen and indirect activation or priming of host immunity (Majumder et al., 2014; Spence et al., 2014). Several isolates recovered in this study inhibited *M. oryzae* in vitro, supporting the potential for direct antagonism. However, differences in host transcriptional responses and chitin-induced ROS production also indicate that microbiome colonization influenced host immune state.

Axenic plants exhibited elevated basal expression of genes associated with both SA- and JA/ET-mediated defense pathways, yet this heightened transcriptional state did not translate into improved disease resistance or enhanced chitin-induced ROS production. Instead, AX plants developed larger lesions, exhibited increased chlorosis, and were substantially more susceptible to blast disease despite expressing numerous defense-associated genes at higher levels than HO plants. These observations indicate that constitutive activation of defense-associated transcriptional programs does not necessarily translate into effective disease resistance. Similar constitutive activation of immune-related genes has been reported in axenically grown rice and has been attributed to the absence of microbial signals that normally regulate immune development and homeostasis (Santos-Medellín et al., 2022).

One interpretation is that microbiome colonization contributes to immune homeostasis rather than simply increasing defense activation. Previous studies have shown that plant-associated microbial communities influence the development and calibration of host immune responses, promoting immune responsiveness while limiting inappropriate or energetically costly defense activation (Santos-Medellín et al., 2022; Teixeira et al., 2021; Yu et al., 2019). Under this framework, the elevated defense-gene expression observed in AX plants may reflect immune dysregulation resulting from the absence of microbial signals that normally help shape immune function. In contrast, microbiome-colonized plants maintained lower basal expression of multiple defense-associated genes while exhibiting greater resistance to blast disease.

The ROS assays further support this interpretation. Although AX plants displayed elevated basal expression of numerous defense-associated genes, HO plants generated significantly stronger ROS bursts following chitin elicitation. ROS production represents one of the earliest outputs of pattern-triggered immunity and contributes to defense signaling, cell wall reinforcement, and restriction of pathogen growth (Bigeard et al., 2015; Gao et al., 2020; Yang et al., 2017).Thus, microbiome colonization may promote effective immune responsiveness and regulation rather than constitutive defense activation. Collectively, the stronger ROS response, reduced disease severity, and more restricted symptom development observed in HO plants suggest that microbiome colonization enhances immune competence while maintaining immune homeostasis. This relationship suggests that the quality of the immune response may be more important than the magnitude of defense-gene expression in determining disease outcome.

An exception to the broader pattern was *OsJAR1*, a key component of jasmonate signaling that catalyzes the formation of the bioactive hormone JA-Ile. Basal *OsJAR1* expression was lower in HO plants than in AX plants despite the greater disease resistance observed in microbiome-colonized plants. One possibility is that reduced constitutive *OsJAR1* expression reduces the costs of sustained jasmonate signaling while preserving the ability to activate jasmonate-dependent defenses when required. Consistent with this interpretation, *OsJAR1* expression increased following pathogen challenge, suggesting that microbiome colonization may alter the regulation of jasmonate signaling rather than simply suppressing it (Shimizu et al., 2013).

The reduced basal expression of several defense-associated genes in HO plants, together with enhanced ROS responsiveness and reduced disease severity, is also broadly consistent with features commonly associated with induced systemic resistance (ISR), which is frequently linked to JA/ET signaling (Kour et al., 2024; Spence et al., 2014; van Wees et al., 1999; Yang et al., 2017). However, the present study did not directly test ISR or immune priming, and alternative explanations, including microbiome-mediated regulation of immune homeostasis, remain possible. Future work examining the kinetics and magnitude of defense activation following pathogen challenge will be required to distinguish between these models.

Interestingly, the strongest transcriptional responses following infection were observed in AXINF plants. Genes associated with oryzalexin S biosynthesis, including *OsDXS3*, *OsCyc1*, and *OsDTC2*, as well as the defense-associated genes *OsPR1A*, *OsPR10B*, and *OsWRKY77*, were induced to substantially higher levels in AXINF plants than in HOINF plants. However, AXINF plants also exhibited the greatest disease severity. These observations suggest that strong induction of defense-associated genes alone does not necessarily equate to effective resistance and may instead reflect a compensatory response to increased pathogen colonization. Consequently, elevated transcriptional activation alone may be a poor predictor of disease outcome. The specific downstream mechanisms responsible for disease restriction in microbiome-colonized plants remain unresolved. Future work should determine whether additional defense outputs, including lignification, callose deposition, or antimicrobial metabolite production, contribute to the enhanced disease resistance observed in HO plants (Gao et al., 2020; Krishnappa et al., 2024; Meena et al., 2020; Velmurugan et al., 2023).

Microbiome colonization, therefore, enhanced blast resistance through effects on host immune regulation and potentially through direct microbial antagonism. Because plant-associated microbial communities can also be reshaped during pathogen challenge, we next examined whether blast infection altered bacterial community structure (Jobert et al., 2025; Liu et al., 2021; Tharp et al., 2026). Axenic plants yielded substantially fewer non-organelle bacterial reads and contained limited representation of several plant-associated bacterial lineages compared with colonized plants. Although amplicon sequencing cannot establish absolute sterility, these data support a strong reduction in bacterial colonization and are consistent with the altered immune phenotype observed in axenic plants.

Blast infection did not significantly alter alpha diversity but did shift beta diversity, indicating that infection changed the relative abundances of existing community members without causing broad loss of richness or evenness. Similar patterns have been observed in other studies of rice-associated microbiomes during disease (Dastogeer et al., 2022; Jobert et al., 2025). This restructuring is compatible with the broader “cry-for-help” model, in which stressed plants modify their interactions with associated microbes in ways that favor potentially beneficial populations (Dastogeer et al., 2022; Jobert et al., 2025; Paasch and He, 2021; Tharp et al., 2026). However, pathogen-driven disruption, altered host physiology, or changes in resource availability could also generate similar compositional shifts. The present data therefore demonstrate infection-associated restructuring but do not distinguish among these possible mechanisms.

Differential abundance analysis identified several taxa that responded to infection. The most strongly enriched ASV was assigned to *Brevibacillus*, a genus containing strains with documented biological control and plant growth-promoting activity (Kim et al., 2024; Liu et al., 2024; Yang et al., 2023). The strong enrichment of the *Brevibacillus*-associated ASV25 is particularly noteworthy because it was the most strongly infection-associated feature detected in the dataset, suggesting that members of this lineage may play an important role in microbiome restructuring during rice blast infection. Notably, the enrichment of *Brevibacillus* occurred despite the absence of major changes in alpha diversity, indicating that infection-associated restructuring was driven by specific taxa rather than broad shifts in community diversity. Infection-enriched ASVs were also assigned to genera such as *Rhodococcus* and *Azospirillum*, which likewise include strains associated with plant growth promotion or disease suppression (Afordoanyi et al., 2023; Lemanceau et al., 2017; Valette et al., 2020b). In contrast, ASVs assigned to broad biological control-associated genera such as *Bacillus* and *Pseudomonas* displayed heterogeneous responses, with closely related ASVs increasing or decreasing following infection.

This variation indicates that infection-associated selection did not occur uniformly at the genus level. Because species-level identification was not reliable at the sequencing resolution used here, these ASVs may represent distinct species, strain-level variants, or closely related lineages with different functional capacities. Plant hosts can influence microbiome assembly at fine taxonomic scales through immune recognition of microbial features, including variation in MAMP epitopes, and these effects may occur below the resolution of conventional 16S rRNA gene surveys (Hacquard et al., 2017). The contrasting responses of closely related ASVs may therefore reflect selection for specific traits or functions rather than broad enrichment of entire genera.

To examine whether compositional differences were associated with microbial function, endophytic isolates were recovered from healthy and infected plants and evaluated for antagonism against *M. oryzae*, plant hormone production, nitrogen fixation, and phosphate solubilization. Of the tested traits, only phosphate solubilization differed significantly by isolation source, with phosphate-solubilizing bacteria recovered more frequently from infected plants. Phosphate availability influences carbohydrate metabolism, redox regulation, signaling, and multiple components of plant defense (Sun et al., 2026). It also has a complex relationship with rice blast, as elevated phosphate accumulation in rice tissue can enhance *M. oryzae* pathogenicity, and the pathogen can manipulate host phosphate-starvation responses to promote phosphate acquisition (Campos-Soriano et al., 2020; Jabbar et al., 2026; Martín-Cardoso et al., 2024). The enrichment of phosphate-solubilizing isolates may therefore indicate infection-associated changes in microbial phosphate cycling. Some phosphate-solubilizing *Pseudomonas* and *Bacillus* strains can also accumulate phosphate intracellularly, raising the possibility that enriched isolates affect phosphate availability within plant tissues (Yang et al., 2025). However, phosphate solubilization and phosphate accumulation were not directly linked in this study, and the consequences of these activities for blast resistance remain unclear. Determining whether infected plants selectively recruit phosphate-transforming bacteria, or whether these isolates alter host or pathogen phosphate availability, will require direct measurements of colonization, phosphate accumulation, and disease effects.

Taxonomic identification of cultured isolates supported differences between communities recovered from healthy and infected plants, but species-level assignments should be interpreted cautiously. MALDI-TOF identified several isolates as members of *Bacillus* and *Pseudomonas*, although colony morphology suggested that some species-level assignments were unreliable. This limitation is particularly relevant within closely related groups such as the *Bacillus cereus* species complex, where discriminatory power can be limited even with genome-based approaches (Liu et al., 2018). Whole-genome sequencing will therefore be necessary to resolve isolate identity and to identify biosynthetic or functional genes that may contribute to disease suppression.

The strongest functional evidence for microbiome-mediated protection came from the SynCom experiments. Two 10-member communities assembled from isolates recovered from healthy or infected plants significantly reduced blast disease severity relative to HOINF plants and reproduced much of the disease-suppressive effect associated with microbiome colonization. These findings demonstrate that disease suppression does not require the full complexity of the soil-derived microbiome and can be recapitulated by a simplified, culturable subset of the rice-associated community. Simplified SynComs have similarly been shown to suppress disease in other plant systems while offering greater reproducibility and experimental tractability than complex natural communities (Li et al., 2021; Liu et al., 2019; Xu et al., 2025).

The SynCom derived from healthy plants produced a greater reduction in disease severity than the community derived from infected plants, although the difference between the two communities was small and not statistically significant. Thus, the greater prevalence of phosphate-solubilizing isolates in the infected-derived community did not translate into superior disease suppression under the conditions tested. This result does not rule out a role for phosphate cycling, but it indicates that phosphate solubilization alone was insufficient to explain SynCom performance. The SynCom experiments suggest that disease suppression likely arises from multiple interacting microbial functions, including immune regulation, direct antagonism, nutrient competition, and the production of antimicrobial metabolites.

The apparent effectiveness of both SynComs also suggests that disease suppression was not dependent on preserving the full complexity of the soil-derived community. However, the underlying community assembly and colonization dynamics remain unknown. Future experiments should quantify the abundance and persistence of individual SynCom members, determine whether all strains are required for protection, and test the communities across rice genotypes, pathogen isolates, and environmental conditions. Genome sequencing and strain-specific tracking would further enable functional prediction and identification of the taxa most responsible for disease suppression.

Collectively, this work establishes a gnotobiotic rice pathosystem for mechanistic investigation of host-microbiome-pathogen interactions and demonstrates that the rice microbiome contributes to blast resistance through coordinated effects on immune regulation, infection-associated community restructuring, and microbial function. Rather than simply increasing defense activation, microbiome colonization was associated with a more effective and responsive immune state characterized by stronger ROS production, reduced disease severity, and lower basal expression of several defense-associated genes. These findings support a model in which the rice microbiome contributes to disease resistance through immune modulation rather than simple activation of defense pathways. By integrating controlled microbiome manipulation, host immune profiling, community sequencing, culture-dependent functional assays, and SynCom reconstruction, this study connects infection-associated microbiome dynamics with experimentally validated disease outcomes and provides a foundation for microbiome-informed strategies for sustainable rice blast management.

## 5. Conclusion

This study establishes a gnotobiotic rice pathosystem for investigating host-microbiome-pathogen interactions and demonstrates that the rice microbiome contributes to blast resistance. Microbiome colonization reduced disease severity, enhanced chitin-induced ROS responsiveness, and was associated with lower basal expression of multiple defense-associated genes, suggesting that effective resistance depends on immune regulation and responsiveness rather than constitutive defense activation alone. Blast infection also altered bacterial community composition, while culture-dependent analyses identified microbial traits associated with disease-associated communities. Furthermore, simplified synthetic communities reproduced much of the disease-suppressive effect observed in microbiome-colonized plants, demonstrating that key protective functions can be retained within defined microbial consortia. Together, these findings provide new insight into microbiome-mediated blast suppression and a foundation for microbiome-informed strategies for sustainable rice disease management.

## CRediT authorship contribution statement

**Jessie Fernandez**: Conceptualization, Methodology, Formal analysis, Validation, Resources, Funding acquisition, Project administration, Supervision, Writing – original draft, Writing – review & editing. **Timothy Johnson**: Conceptualization, Methodology, Investigation, Data curation, Writing – original draft. **Lainey Kermerer**: Investigation, Writing – review & editing. **Nalleli Payne**: Investigation, Writing – review & editing. Matthew VanWeelden: Resources, Writing – review & editing. **Luiz F. Wurdig Roesch**: Formal analysis, Supervision, Writing – review & editing.

## Acknowledgments

We thank the members of the Fernandez Lab for valuable feedback, critical review, and insightful discussions throughout the development of this work and manuscript. We also thank members of the Sheng Yang He Lab at Duke University for their guidance and advice regarding the design and implementation of the gnotobiotic growth system. This work was supported by the University of Florida Institute of Food and Agricultural Sciences (UF/IFAS) through a Research Opportunity Seed Fund (ROSF) award.

## Appendix A. Supplementary material

### Data availability

Data will be made available on request. Raw 16S rRNA amplicon sequencing data have been deposited in the NCBI Sequence Read Archive (SRA) under BioProject accession PRJNA1546142.

