## Supplemental Materials for "A Gnotobiotic Rice Pathosystem Reveals How the Microbiome Suppresses Rice Blast Disease"

**Supplementary Data**

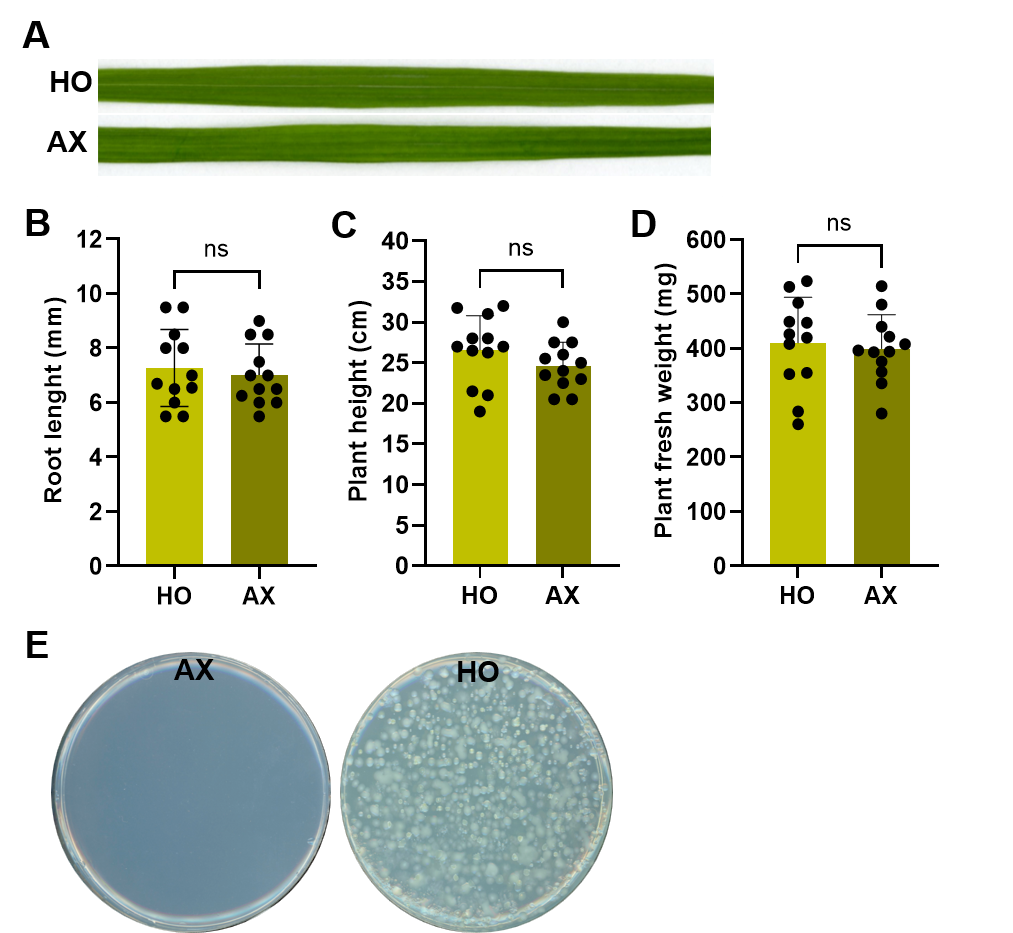

**Figure S1.** Growth and microbial colonization of axenic and holoxenic rice plants in the gnotobiotic system. (A) Representative rice leaves from AX and HO plants grown in the gnotobiotic system. (B–D) Quantification of root length, plant height, and plant fresh weight. Data are presented as mean ± SD, with individual biological replicates shown as dots. Each experiment included four plants per treatment and was independently repeated three times (total plants = 12; *n* = 3). Statistical differences between AX and HO plants were assessed using Welch’s unpaired *t*-test; ns, not significant. (E) Representative R2A plates showing culture-based microbial recovery from surface-sterilized leaves of AX and HO plants. Representative results from three independent experiments are shown.

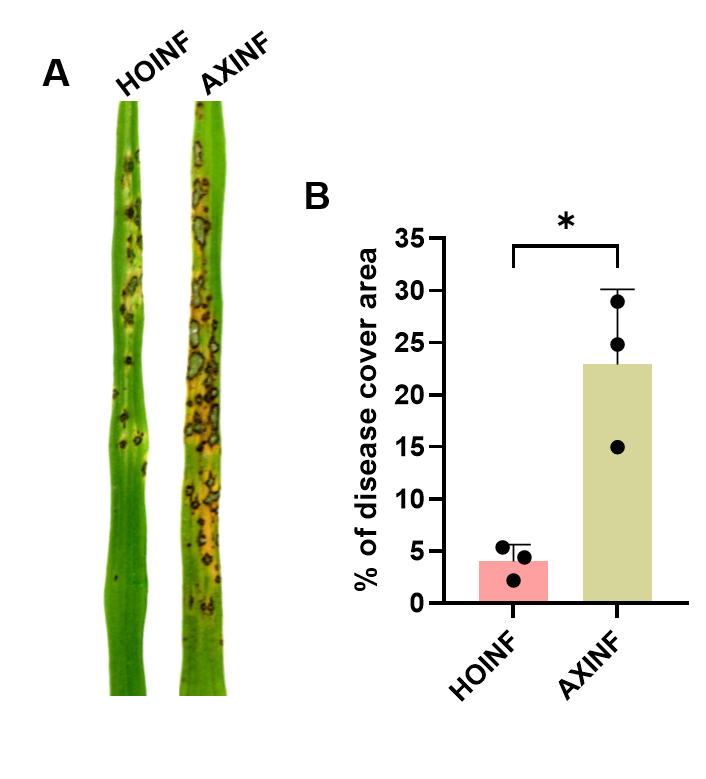

**Figure S2**. Holoxenic rice plants exhibit reduced blast disease severity compared with axenic plants. (A) Representative leaves of the rice cultivar YT16 showing blast symptoms following spray inoculation with *M. oryzae*. (B) Disease severity quantified as percentage disease-covered leaf area. Bars represent mean ± SD. Statistical significance was determined using Welch's *t*-test. * indicates *P* < 0.05.

**Table S1.** Distribution of raw and retained sequencing reads following quality control and filtering.

| Sample | Treatment | Batch | Raw Reads | Reads After Organelle Filtering |
| --- | --- | --- | --- | --- |
| zr28470_1V3V4 | HO | 1 | 96218 | 434 |
| zr28470_2V3V4 | HO | 1 | 78890 | 1147 |
| zr28470_3V3V4 | HO | 1 | 86831 | 865 |
| zr28470_4V3V4 | HOINF | 1 | 85690 | 14086 |
| zr28470_5V3V4 | HOINF | 1 | 79493 | 30065 |
| zr28470_6V3V4 | HOINF | 1 | 76305 | 39564 |
| zr28470_7V3V4 | AX | 1 | 95182 | 1694 |
| zr28470_8V3V4 | AX | 1 | 73037 | 262 |
| zr28470_9V3V4 | AX | 1 | 98071 | 7735 |
| zr29672_1V3V4 | HO | 2 | 89574 | 3300 |
| zr29672_2V3V4 | HO | 2 | 111978 | 1613 |
| zr29672_3V3V4 | HO | 2 | 104515 | 1192 |
| zr29672_4V3V4 | HOINF | 2 | 102536 | 647 |
| zr29672_5V3V4 | HOINF | 2 | 87367 | 93 |
| zr29672_6V3V4 | HOINF | 2 | 113469 | 63 |
| zr29672_7V3V4 | AX | 2 | 97051 | 196 |
| zr29672_8V3V4 | AX | 2 | 102919 | 918 |
| zr29672_9V3V4 | AX | 2 | 115712 | 89 |
| zr30737_1V3V4 | HO | 3 | 36280 | 3218 |
| zr30737_2V3V4 | HO | 3 | 45861 | 3196 |
| zr30737_3V3V4 | HO | 3 | 72990 | 2554 |
| zr30737_4V3V4 | HOINF | 3 | 37209 | 16320 |
| zr30737_5V3V4 | HOINF | 3 | 42145 | 5391 |
| zr30737_6V3V4 | HOINF | 3 | 51986 | 9363 |

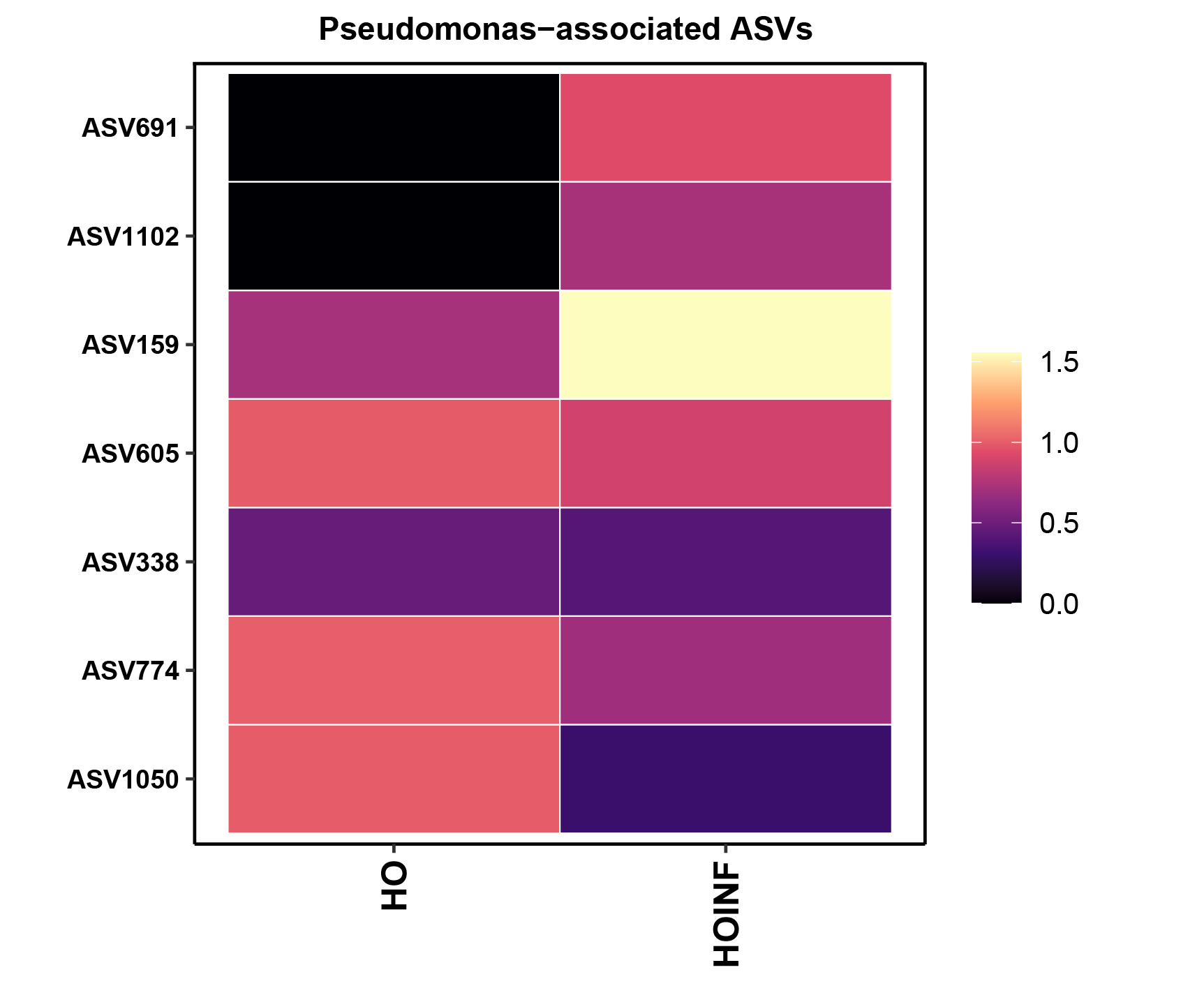

**Figure S3.** Heatmap of significantly differentially abundant ASVs assigned to the genus *Pseudomonas*. Heatmap showing DESeq2-normalized counts summarized by treatment and transformed as log10(normalized counts + 1) of differentially abundant *Pseudomonas* ASVs in HO and HOINF samples. Rows represent individual ASVs and columns represent treatment groups. The analysis was performed on the filtered, unrarefied dataset after removing samples with fewer than 1,000 non-organelle bacterial reads.

**Table S2**. List of PCR primers used for qRT-PCR.

| **Names** | **Primer Sequences (5’-3’)** | **Source** |
| --- | --- | --- |
| EDS1 | Forward: TTGAATTTTGTCGTGCCAGTAGA  Reverse: GGCAGATGCAAGCGGAGTAA | (Patel et al., 2022) |
| WRKY45 | Forward: CGGGTAAAACGATCGAAAGA  Reverse: TTTCGAAAGCGGAAGAACAG | (Akagi et al., 2014) |
| EIL1 | Forward: ACAATGCCACGATCATGGAG  Reverse: TCAGTAGTACCAATTCGAGC | (Ke et al., 2014a) |
| EIN2 | Forward: CAAGGAACCAGTGACAACCA  Reverse: GCAGTCGTCTCCGCAGTTAG | (Duan et al., 2014) |
| PR10b | Forward: AGGTGTTGGAGGTTAAGAGC  Reverse: TCGACGATCATCTTCTCCTG | (Valette et al., 2020a) |
| ERF1 | Forward: CATATCACCTTGACGCCCCA  Reverse: ACCCTCACAAACTCACTCGG | (Spence et al., 2014) |
| WRKY77 | Forward: CTGTGTCCAGCTACCTCTCC  Reverse: TGAAGAGAGCGATCACCTC | (Spence et al., 2014) |
| JAR1 | Forward: TCTCCCCAGCCTTAACCGTA  Reverse: CTAAACGCGACGACAAACCC | (Spence et al., 2014) |
| PAL | Forward: GCACATCTTGGAGGGAAGCT  Reverse: GCGCGGATAACCTCAATTTG | (Valette et al., 2020b) |
| DXS3 | Forward: TGTTCTTGCCAGACAGGTAC  Reverse: GTCGGCTGATGTGTATATGC | (Valette et al., 2020b) |
| CYC1 | Forward: CCATCTGCTGCAAACTTTCC  Reverse: TCCTGCATCTCCAAATCCAC | (Valette et al., 2020b) |
| DTC2 | Forward: AGAGAATATCGGTTCCTGGTC  Reverse: CCGTTTACGTTGAATCGCT | (Valette et al., 2020b) |
| PR1A | Forward: CGTCTTCATCACCTGCAACTACTC  Reverse: CATGCATAAACACGTAGCATAGC | (Ke et al., 2014b) |
| WRKY62 | Forward: GAGACCGTGAAGGATGGG  Reverse: TCGTAGGTCGCCACTAGC | (Valette et al., 2020b) |
| GAPDH | Forward: AAGCCAGCATCCTATGATCAGATT  Reverse: CGTAACCCAGAATACCCTTGAGTTT | (Chen et al., 2020) |
